# Bayesian Evaluation of Temporal Signal in Measurably Evolving Populations

**DOI:** 10.1101/810697

**Authors:** Sebastian Duchene, Philippe Lemey, Tanja Stadler, Simon YW Ho, David A Duchene, Vijaykrishna Dhanasekaran, Guy Baele

## Abstract

Phylogenetic methods can use the sampling times of molecular sequence data to calibrate the molecular clock, enabling the estimation of evolutionary rates and timescales for rapidly evolving pathogens and data sets containing ancient DNA samples. A key aspect of such calibrations is whether a sufficient amount of molecular evolution has occurred over the sampling time window, that is, whether the data can be treated as having come from a measurably evolving population. Here we investigate the performance of a fully Bayesian evaluation of temporal signal (BETS) in sequence data. The method involves comparing the fit to the data of two models: a model in which the data are accompanied by the actual (heterochronous) sampling times, and a model in which the samples are constrained to be contemporaneous (isochronous). We conducted simulations under a wide range of conditions to demonstrate that BETS accurately classifies data sets according to whether they contain temporal signal or not, even when there is substantial among-lineage rate variation. We explore the behaviour of this classification in analyses of five empirical data sets: modern samples of *A/H1N1 influenza virus*, the bacterium *Bordetella pertussis*, coronaviruses from mammalian hosts, ancient DNA from *Hepatitis B virus* and mitochondrial genomes of dog species. Our results indicate that BETS is an effective alternative to other tests of temporal signal. In particular, this method has the key advantage of allowing a coherent assessment of the entire model, including the molecular clock and tree prior which are essential aspects of Bayesian phylodynamic analyses.

## Introduction

The molecular clock has become a ubiquitous tool for studying evolutionary processes in rapidly evolving organisms and in data sets that include ancient DNA. In its simplest form, the molecular clock posits that evolutionary change occurs at a predictable rate over time (Zuckerkandl and Pauling 1965). The molecular clock can be calibrated to estimate divergence times by using sampling time information, the timing of known divergence events, or a previous estimate of the evolutionary rate (Hipsley and Müller 2014). For example, Korber et al. (2000) used sampling times to calibrate the molecular clock and to infer the time of origin of HIV group 1. Their approach consisted of estimating a phylogenetic tree and conducting a regression of the distance from the root to each of the tips as a function of sequence sampling times. In this method, the slope of the regression is an estimate of the evolutionary rate in substitutions per site per unit of time, the intercept with the time axis is the age of the root node, and the coefficient of determination (*R*^2^) is the degree to which the data exhibit clocklike behaviour (Rambaut et al. 2016). Despite the practicality of root-to-tip regression, its use as a statistical tool for molecular dating has several well-known limitations. In particular, data points are not independent because they have shared ancestry (i.e., internal branches are traversed multiple times) and a strict clocklike behaviour is assumed by necessity.

The past few decades have seen a surge in novel molecular clock models that explicitly use phylogenetic information. Bayesian methods have gained substantial popularity, largely due to the wide array of complex models that can be implemented and the fact that independent information, including calibrations, can be specified via prior distributions (Huelsenbeck et al. 2001; Nascimento et al. 2017). Of particular importance is the availability of molecular clock models that relax the assumption of a strict clock by explicitly modelling rate variation among lineages (reviewed by Ho and Duchene (2014) and by Bromham et al. (2018)).

Regardless of the methodology used to analyse time-stamped sequence data, a sufficient amount of molecular evolution must have occurred over the sampling time window to warrant the use of sequence sampling times for calibration. In such cases, the population can be considered to be ‘measurably evolving’ (Drummond et al. 2003). The degree of ‘temporal information’ in sequence data is determined by the sequence length, the evolutionary rate, the range of available sampling times, and the number of sequences. Some viruses evolve at a rate of around 5×10^−3^ subs/site/year (Duchene et al. 2014), such that samples collected over a few weeks can be sufficient to calibrate the molecular clock. In more slowly evolving organisms, such as mammals, a sampling window of tens of thousands of years might be necessary; this can be achieved by including ancient DNA sequences (Drummond et al. 2003; Biek et al. 2015).

Testing for temporal signal is an important step prior to interpreting evolutionary rate estimates (Rieux and Balloux 2016). A data set is considered to have temporal signal if it can be treated as a measurably evolving population, defined by Drummond et al. (2003) as “populations from which molecular sequences can be taken at different points in time, among which there are a statistically significant number of genetic differences”. In general, the presence of temporal signal also implies that the data set will produce reliable divergence time estimates (Murray et al. 2015). A popular method to assess temporal signal is the date-randomization test that compares actual evolutionary rate estimates to those obtained by repeatedly permuting the sequence sampling times (Ramsden et al. 2009). A data set is considered to have strong temporal signal if the rate estimated using the correct sampling times does not overlap with those of the permutation replicates (Duchêne et al. 2015; Murray et al. 2015; Duchene et al. 2018). An implementation of this test is also available that performs the permutation during a single Bayesian analysis (Trovão et al. 2015). The interpretation of the date-randomization test is essentially frequentist in nature, which leads to an inconsistent mixture of statistical frameworks when Bayesian phylogenetic methods are used. Moreover, the procedure is not applicable in cases with small numbers of sampling times, owing to the limited number of possible permutations (Duchêne et al. 2015).

We propose a fully Bayesian model test, which we refer to as BETS (Bayesian Evaluation of Temporal Signal), to assess temporal signal based on previous analyses by Baele et al. (2012). The approach involves quantifying statistical support for two competing models: a model in which the data are accompanied by the actual sampling times (i.e., the data are treated as heterochronous) and a model in which the sampling times are contemporaneous (i.e., the data are treated as isochronous). Therefore, the sampling times are treated as part of the model and the test can be understood as a test of ultrametricity of the phylogenetic tree. If incorporating sampling times improves the statistical fit, then their use for clock calibration is warranted. The crux of BETS, as with Bayesian model selection, is that it requires calculating the marginal likelihood of the model in question. The marginal likelihood measures the evidence for a model given the data, and calculating it requires integration of its likelihood across all parameter values, weighted by the prior (Kass and Raftery 1995).

Because the marginal likelihood is a measure of model evidence, the ratio of the marginal likelihoods of two competing models, known as the Bayes factor, is used to assess support for one model relative to the other. In the case of applying BETS, let *M*_het_ represent the heterochronous model, *M*_iso_ the isochronous model, and *Y* the sequence data, such that P(*Y*|*M*_het_) and P(*Y*|*M*_iso_) are their respective marginal likelihoods. These models differ in the number of parameters. In *M*_iso_ the evolutionary rates and times are nonidentifiable, so the rate is fixed to an arbitrary value; in *M*_het_ the rate is a free parameter. Differences in the number of parameters do not need to be taken into account separately, because accurate marginal likelihood estimators naturally penalize excessive parameterization. Kass and Raftery (1995) provide guidelines for interpreting Bayes factors, where a (log) Bayes factor log(P(*Y*|*M*_het_)) – log(P(*Y*|*M*_iso_)) of at least 5 indicates ‘very strong’ support for *M*_het_ over *M*_iso_, a value of 3 indicates ‘strong’ support, and a value of 1 is considered as positive evidence for *M*_het_ over *M*_iso_.

The importance of model selection in Bayesian phylogenetics has prompted the development of various techniques to calculate log marginal likelihoods (reviewed by Baele et al. (2014) and by Oaks et al. (2019)). These techniques can be broadly classified into prior-based and/or posterior-based estimators and path sampling approaches. Prior- and posterior-based estimators, also known as importance sampling, include the widely used harmonic mean estimator (Newton and Raftery 1994) and the AICM and BICM (Bayesian analogues to the Akaike information criterion and the Bayesian information criterion, respectively) (Raftery et al. 2007). These scores are easy to compute because they only require samples from the posterior distribution as obtained through Markov chain Monte Carlo (MCMC) integration. However, the harmonic mean estimator has been shown to have unacceptably high variance when the prior is diffuse relative to the posterior, and, together with the AICM, has shown poor performance in practical settings (Baele et al. 2012, 2013). The BICM requires a sample size to be specified for each parameter, which is far from trivial for phylogenetic inference and therefore remains unexplored for such applications.

Path sampling approaches include path sampling (originally introduced in phylogenetics as ‘thermodynamic integration’) (Lartillot and Philippe 2006), stepping-stone sampling (Xie et al. 2011), and generalized stepping-stone (GSS) sampling (Fan et al. 2011; Baele et al. 2016). These methods depend on drawing samples using MCMC from a range of power posterior distributions that represent the path from the posterior to the (working) prior, and therefore require additional computation. Another numerical technique that was recently introduced to phylogenetics is nested sampling (NS) (Maturana et al. 2019), which approximates the log marginal likelihood by simplifying the marginal-likelihood function from a multi-dimensional to a one-dimensional integral over the cumulative distribution function of the log marginal likelihood (Skilling 2006). Fourment et al. (2020) recently compared the accuracy of a range of methods for estimating log marginal likelihoods and found GSS to be the most accurate, albeit at increased computational cost. Clearly, the reliability of the log marginal likelihood estimator is a key consideration for applying BETS.

We conducted a simulation study to assess the reliability of BETS under a range of conditions that are typical for data sets of rapidly evolving organisms and of those that include ancient DNA. We also analysed five empirical data sets to showcase the performance of the test in practice. Our analyses demonstrate the utility of BETS in providing accurate evaluation of temporal signal across a wide range of situations.

## Results

### Simulations of Measurably Evolving Populations

In our simulations we considered sequence data from heterochronous and isochronous trees. Heterochronous trees represent a situation where there is sufficient temporal signal, whereas isochronous trees lack temporal signal altogether. We simulated heterochronous phylogenetic trees under a stochastic birth-death process with between 90 and 110 tips (fig. 1A and 1B). To generate isochronous trees we used similar settings, but we assumed a single sampling time (fig. 1C). We then simulated evolutionary rates along the trees according to an uncorrelated relaxed clock with an underlying lognormal distribution with a mean of 5×10^−3^ subs/site/unit time and a standard deviation, *σ*, of 0.0, 0.1, 0.5, or 1, where *σ*=0.0 is equivalent to simulating under a strict clock. We then simulated sequence evolution using an HKY+Γ substitution model, with parameter values similar to those estimated for influenza virus (Hedge et al. 2013), to generate alignments of 4,000 nucleotides.

**FIG. 1.**
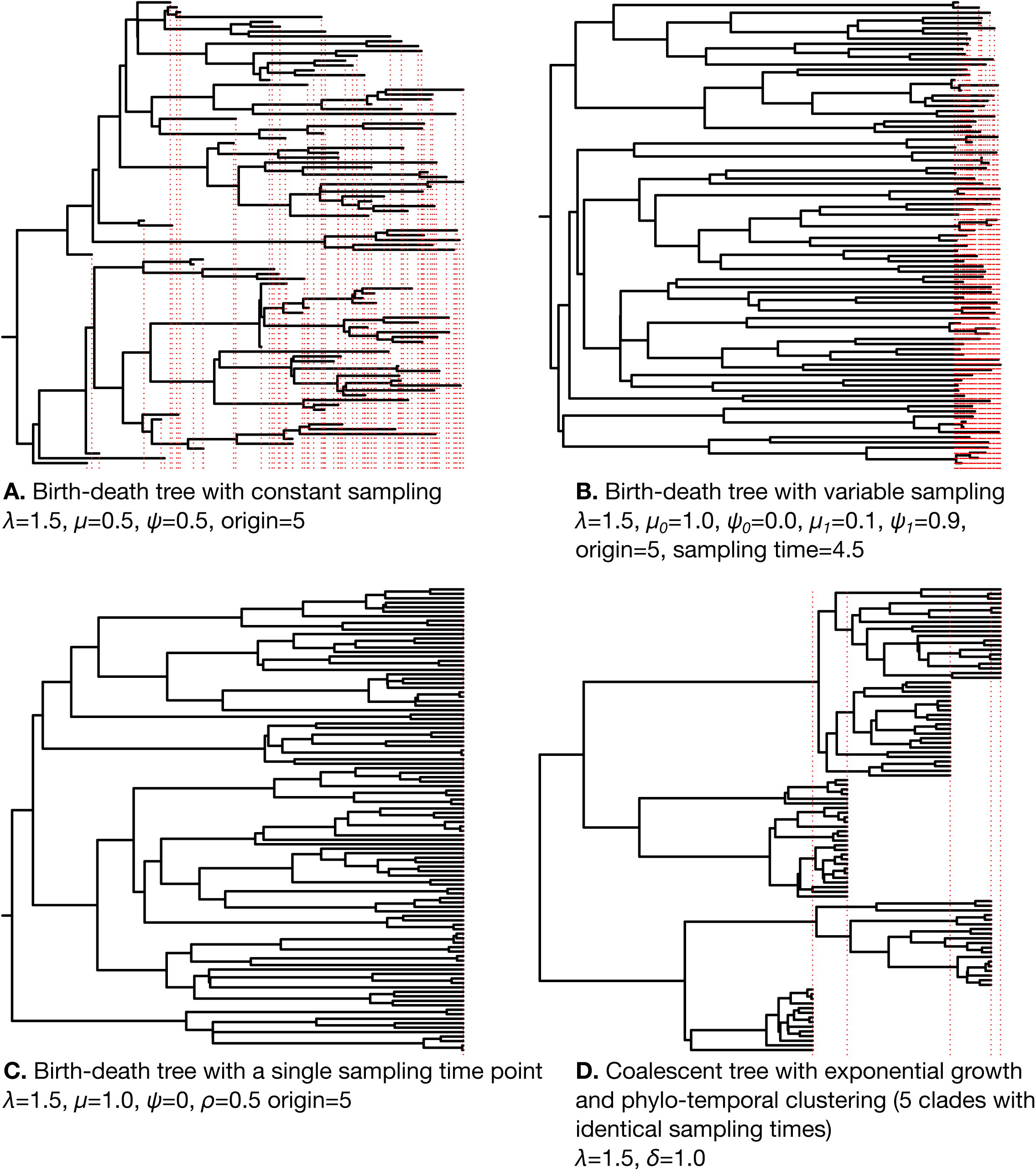
Four examples of phylogenetic trees used in simulations. Red dashed vertical lines indicate the times of the tips and therefore represent the sampling process over time. Trees A–C were simulated under a birth-death process with time of origin of 5, such that the sum of the tree height and the length of the stem branch leading to the root is always 5. Tree D was generated under a coalescent process with exponential growth. The coalescent and birth-death models have an exponential growth rate, *r*, defined as the difference between the birth rate,*λ*, and the become-uninfectious rate,*δ*, such that *r = λ - δ*. We set *λ*=1.5, and *δ*=1. In the birth-death model *δ*=*μ*+*ψ*, where *μ* is the death rate and *ψ* is the sampling rate upon death. Thus, the population growth rate is constant and the same across all trees. Tree A represents a constant sampling process and a wide sampling window (*ψ*=0.5 time units throughout the whole process), whereas in tree B sampling starts after 4.5 time units. Before this time the sampling rate, *ψ*_0_, is zero. After 4.5 time units the sampling rate *ψ*_1_ is 0.9 (and thus *μ*_*1*_= 0.1), resulting in a narrow sampling window. Tree C has samples drawn at a single point in time with a sampling probability at present, *ρ*, of 0.5 (and thus *ψ*=0). Tree D represents a situation where tips with identical sampling times form monophyletic groups, a pattern known as phylo-temporal clustering. To generate these conditions, we used a coalescent model conditioned on the number of tips and their sampling times. These sampling times corresponded to 5 quantiles of a birth-death process with the same *r*.

Our main simulation conditions produced data sets in which about 50% of the sites were variable. We refer to this simulation scenario as (i) ‘high evolutionary rate and wide sampling window’, and we considered three other simulation scenarios that involved (ii) a lower evolutionary rate of 10^−5^ subs/site/unit time, (iii) a narrower sampling window, and (iv) both of the previous two conditions. For a subset of conditions, we investigated the effect of phylo-temporal clustering, a situation in which sequences have been sampled at only a few specific time points and form monophyletic groups (fig. 1D). This pattern has been shown to be a confounding factor that misleads date-randomization tests of temporal signal and that often produces biased evolutionary rate estimates (Duchêne et al. 2015; Murray et al. 2015; Tong et al. 2018).

We analysed the sequence data using a strict clock and an uncorrelated relaxed clock with an underlying lognormal distribution (Drummond et al. 2006). We considered three configurations for sampling times: birth-death sampling times, which are correct for the heterochronous data but not for the isochronous data; identical sampling times, which is correct for isochronous data but not for the heterochronous data; and permuted birth-death sampling times, which are incorrect for both heterochronous and isochronous data. We estimated the log marginal likelihoods of these six combinations of sampling times and clock models using NS and GSS as implemented in BEAST 2.5 (Bouckaert et al. 2019) and BEAST 1.10 (Suchard et al. 2018), respectively. Our BETS approach ranked the models according to their log marginal likelihoods and computed log Bayes factors of the best relative to the second-best model and of the best heterochronous model (*M*_het_) compared with the best isochronous model (*M*_iso_).

#### (i) Simulations with High Evolutionary Rate and Wide Sampling Window

Both NS and GSS correctly classified data sets as being heterochronous or isochronous in 10 out of 10 simulations, including in the presence of a high degree of among-lineage rate variation (i.e., *σ*=1.0; figs. 2 and 3 for heterochronous data and supplementary figs. S1 and S2, Supplementary Material online, for isochronous data). Although both log marginal likelihood estimators detected temporal signal, NS supported the relaxed clock over the strict clock for three heterochronous data sets simulated without among-lineage rate variation (*σ*=0.0) and for six data sets simulated with low among-lineage rate variation (*σ* =0.1). In the simulations of isochronous data, NS often favoured the relaxed clock over the strict clock when there was low among-lineage rate variation (*σ*=0.0 and *σ*=0.1), albeit mostly with log Bayes factors below 5 (supplementary fig. S2, Supplementary Material online). In contrast, GSS always selected the strict clock under these conditions (supplementary fig. S1, Supplementary Material online).

**FIG. 2.**
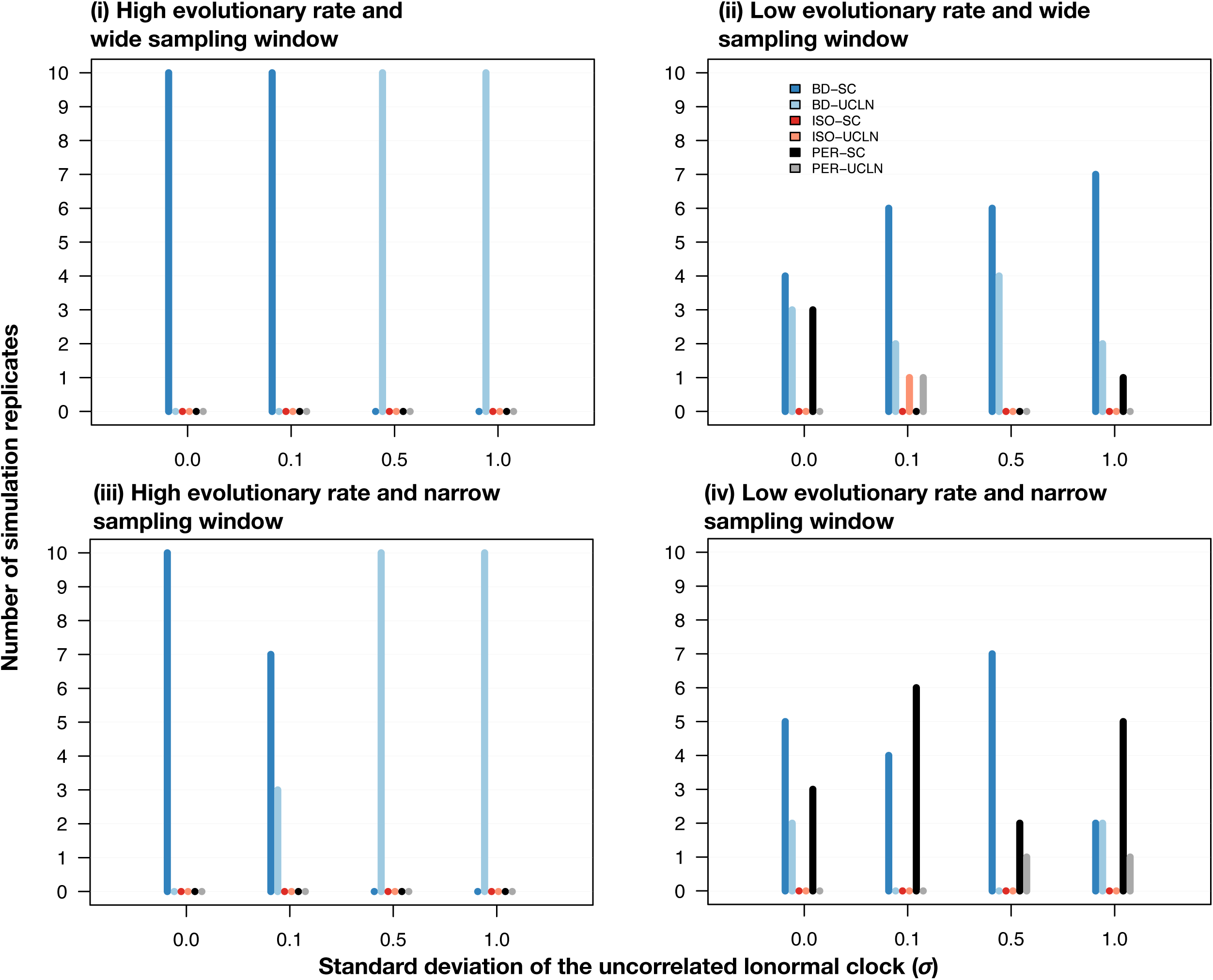
Models selected for heterochronous data using generalized stepping-stone sampling under two evolutionary rates, shown in each panel and noted in the main text as conditions (i) and (ii), and four degrees of among-lineage rate variation as determined by the standard deviation of a lognormal distribution,*σ* (along the *x*-axis). Each set of bars corresponds to a model, with bar heights (along the *y*-axis) representing the number of times each model was selected out of ten simulation replicates. The bars are coloured according to the settings in the analysis, based on combinations of two molecular clock models, strict clock (SC) and the uncorrelated relaxed clock with an underlying lognormal distribution (UCLN), and three settings for sampling times: generated under the birth-death process (BD), identical sampling times (Isochronous; ISO), and permuted (Permuted; PER).

**FIG. 3.**
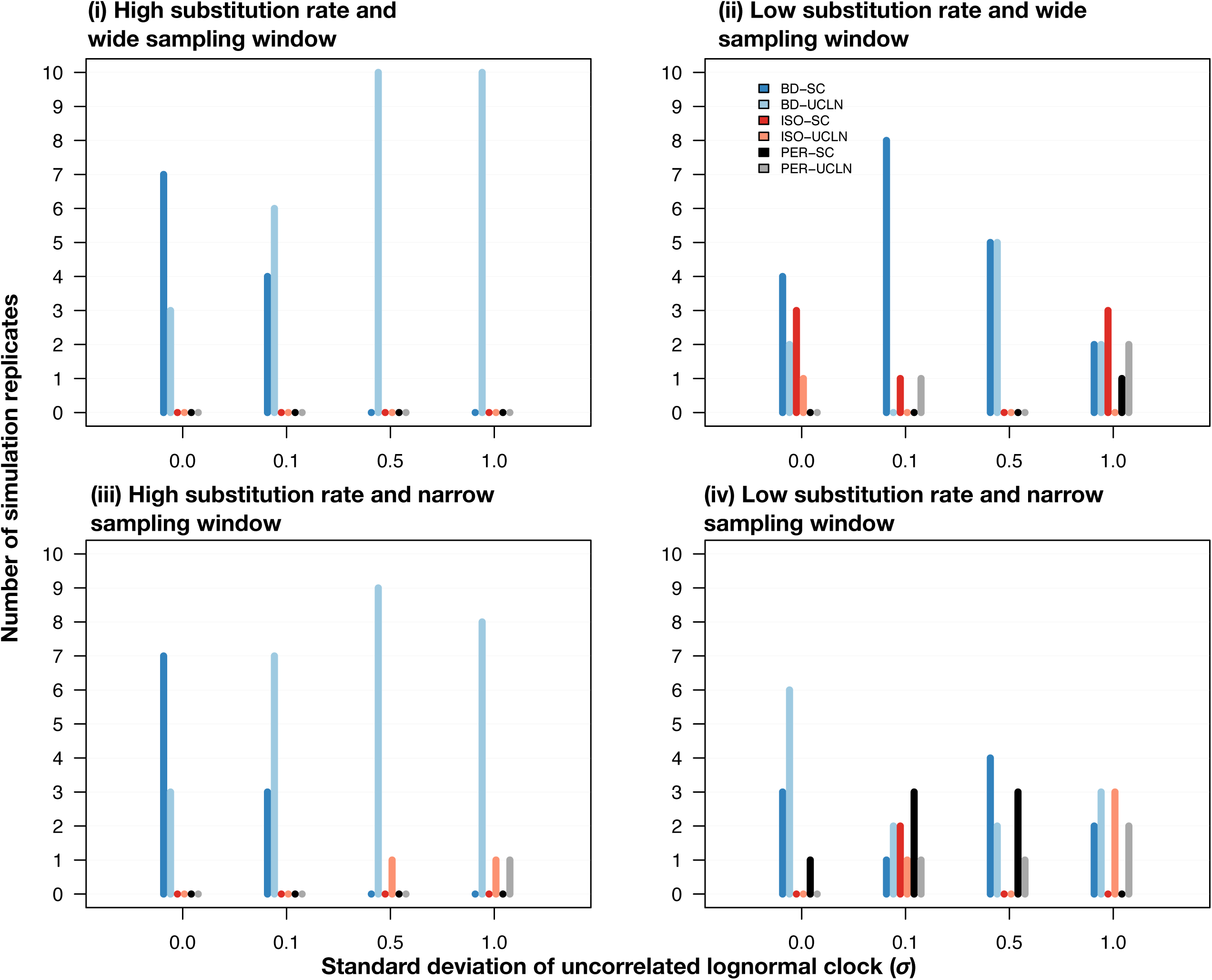
Models selected for heterochronous data using nested sampling under different simulation conditions; four combinations of evolutionary rate and width of the sampling window shown in each panel and noted in the main text as conditions (i) through (iv), and four degrees of among-lineage rate variation as determined by the standard deviation of a lognormal distribution,*σ* (along the *x*-axis). Each set of bars corresponds to a model and their height (along the *y*-axis) represents the number of times each model was selected out of ten simulation replicates. The bars are coloured depending on the analyses settings with two molecular clock models, strict clock (SC) and the uncorrelated relaxed clock with an underlying lognormal distribution (UCLN), and three settings for sampling times: generated under the birth-death process (BD), identical sampling times (Isochronous; ISO), and permuted (Permuted; PER).

For the heterochronous data sets, NS and GSS always displayed very strong support for *M*_het_ over *M*_iso_, with log Bayes factors of at least 90. For the isochronous data sets, the log Bayes factors for *M*_iso_ relative to *M*_het_ were overall much lower, but still decisive, ranging from 30 to 50. Furthermore, log Bayes factors tended to decline with an increasing degree of among-lineage rate variation in the data. Another important observation is that in the heterochronous data, the relaxed clock was consistently selected over the strict clock when assuming that the data were isochronous, or when the sampling times had been permuted (fig. S3, Supplementary Material online). Moreover, the strict clock with permuted sampling times yielded the lowest log marginal likelihoods for heterochronous data. Both of these patterns are likely to be due to an apparently higher degree of among-lineage rate variation when sampling times are misspecified.

#### (ii) Simulations with Low Evolutionary Rate and Wide Sampling Window

Our simulations with a low evolutionary rate of 10^−5^ subs/site/unit time produced data sets that each had on average 10 variable sites (with several replicates only having as few as 4 variable sites), which provides very little information to estimate evolutionary parameters and low power to differentiate between models. Marginal likelihood estimator variance adds to the difficulty in distinguishing between competing models in such conditions. For the heterochronous data sets, GSS selected the heterochronous model with correct dates in at least 7 out of 10 simulation replicates (fig. 2). Across the simulations with different clock models (40 in total), only in five heterochronous data sets did we find models with permuted sampling times to have the highest log marginal likelihoods. For NS, in 12 out of 40 simulations, either isochronous models or those with random sampling times were incorrectly selected when heterochronous data sets were analysed (fig. 3).

Log marginal likelihoods calculated using GSS tended to support models with sampling times (either permuted or those from the birth-death) for the isochronous data, whereas NS appeared to provide equal support for all models (supplementary figs. S1 and S2, Supplementary Material online). However, a critical feature of the results from the data sets with a low evolutionary rate is that the log marginal likelihoods for all models were more similar to one another than those for the data sets with high evolutionary rate (supplementary fig. S4, Supplementary Material online; note that the log marginal likelihood scale in fig. S4 is smaller than that in fig. S3). As a case in point, for the isochronous data with *σ*=0.1 there were log Bayes factors of about 0.1 for the best model with birth-death sampling times relative to those with permuted sampling times. This result points to difficulties distinguishing between models due to estimator variance in the case of few unique site patterns. Additionally, this shows that comparing models with permuted sampling times might be useful for determining whether the data are informative about a particular set of sampling times.

#### (iii) Simulations with High Evolutionary Rate and Narrow Sampling Window

We conducted a set of simulations similar to those described in scenario (i) but where sequence sampling spanned only the last 10% of the age of the tree (0.5 units of time, compared with 5 units of time for the simulations with a wide sampling window; fig. 1B). These conditions reflect those of organisms with deep evolutionary histories and for which samples are available for only a small (recent) portion of this time. Since in these trees the samples were collected over a narrower time window, we used a higher sampling probability to obtain about 100 samples, as in our other simulations. For these analyses we only considered heterochronous data because the isochronous case is identical to the one in scenario (i).

Both GSS and NS showed excellent performance in detecting temporal signal in this scenario, with GSS always selecting models with correct sampling times (fig. 2 and fig. 3). The exceptions to this pattern occurred for one data set with *σ*=0.5 and for two data sets with *σ*=1.0 for NS (fig. 3). Differentiating between the strict clock and relaxed clock appeared somewhat more difficult, particularly for NS, where the relaxed clock with correct sampling times yielded log marginal likelihoods very similar to those for the strict clock for data with low among-lineage rate variation (*σ* of 0.0 or 0.1). Although NS and GSS performed well in these simulations, the log Bayes factors for *M*_het_ relative to *M*_iso_ were much lower than those for data with a high evolutionary rate and a wide sampling window in (i). One obvious example is in the data with *σ*=0.0, where the mean log Bayes factors for *M*_het_ over *M*_iso_ using GSS was 203.15 with a wide sampling window, but decreased to 35.77 when sampling spanned a narrow time window (supplementary fig. S5, Supplementary Material online).

#### (iv) Simulations with Low Evolutionary Rate and Narrow Sampling Window

We considered data sets with a narrow sampling window, as in scenario (iii), and with a low evolutionary rate of 10^−5^ subs/site/unit time, as in scenario (ii). We generated only heterochronous trees under these conditions, because the isochronous case would be identical to (ii).

Estimates of log marginal likelihoods with GSS and NS were very similar among models, with mean log Bayes factors among data sets of less than 1 for the two models with highest log marginal likelihoods for GSS (supplementary fig. S6, Supplementary Material online). In the data sets with *σ*=0.0, GSS and NS always preferred a heterochronous model. However, in a few cases (three for GSS and one for NS) the model with permuted sampling times was selected, indicating that temporal signal was not detected (figs. 2 and 3). As with the data sets with low evolutionary rate and constant sampling (ii), the relaxed clock was occasionally preferred over the strict clock, even when the data sets had no rate variation among lineages.

##### Accuracy of evolutionary rate estimates

We compared the accuracy and precision in rate estimates for our heterochronous simulations with conditions (i) through (iv) using the correct sampling times and the strict and uncorrelated relaxed lognormal clock models. In data sets simulated under a high evolutionary rate and wide sampling window, i.e. condition (i), analyses of all simulation replicates with *σ*=0.0 and *σ*=0.1 had 95% highest posterior density (HPD) intervals that included the true value of the clock rate used to generate the data, 5×10^−3^ subs/site/unit time (fig. 4). When *σ*=0.5, the accuracy was lower, with four data sets analysed under the strict clock and three under the relaxed clock with 95% HPD intervals that included the true value. With *σ*=1.0, only one replicate using the strict clock included this true value in its HPD interval. Importantly, however, under these simulation conditions the HPD intervals of all estimates were within the 95-percentile width of a lognormal distribution with mean 5×10^−3^ and *σ*=0.1 or 0.5 (fig. 4), such that they overlap the evolutionary rate distribution used to generate the data.

**FIG. 4.**
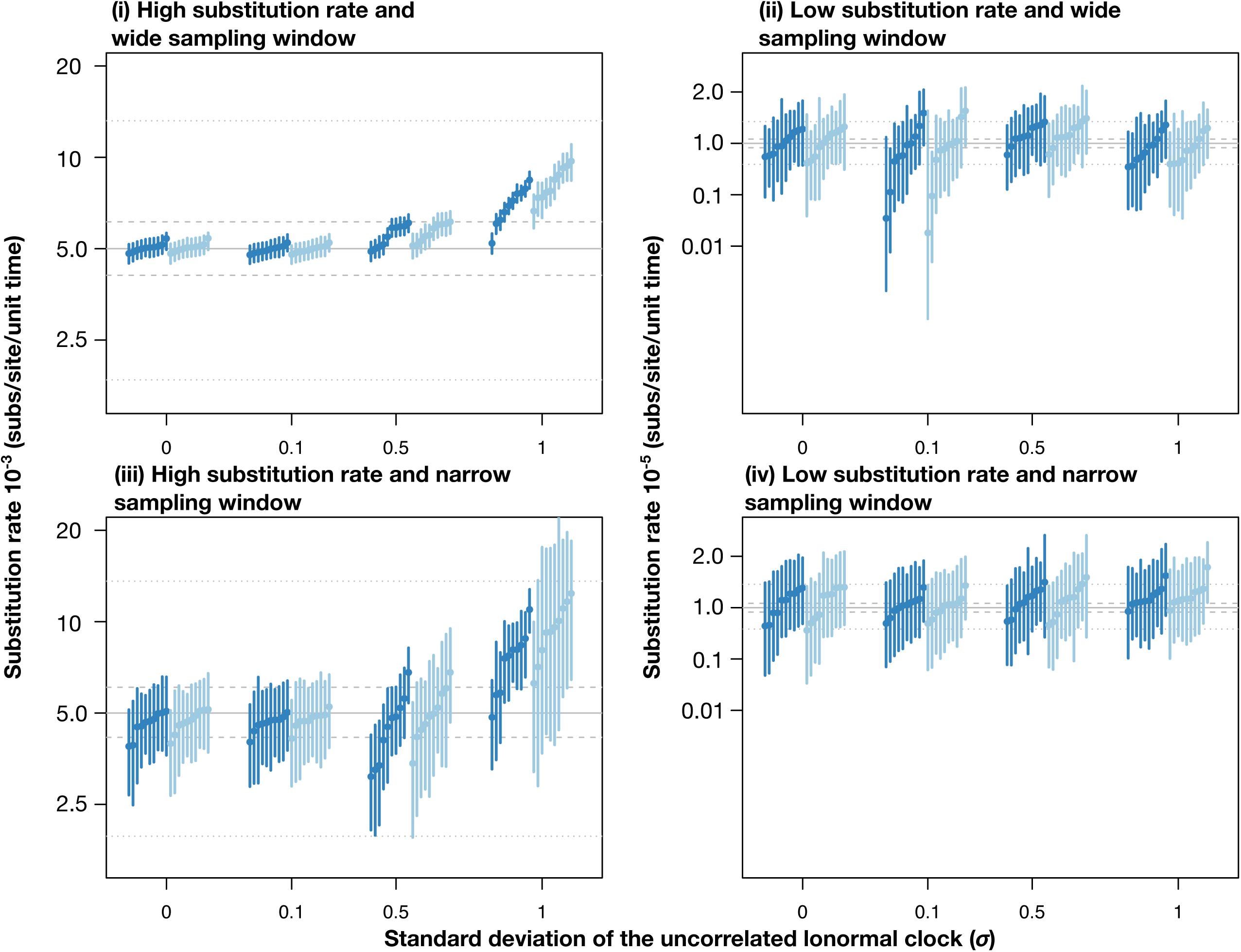
Evolutionary rate estimates for heterochronous data with correct sampling times using a strict clock (in dark blue) and an uncorrelated relaxed clock with an underlying lognormal distribution (in light blue). The panels correspond to the simulation conditions (i) through (iv), described in the main text. The *x*-axis denotes four degrees of among-lineage rate variation used to generate the data, as determined by the standard deviation of a lognormal distribution,*σ*. The *y*-axis corresponds to the evolutionary rate estimate. Solid grey lines correspond to the mean evolutionary rate value used to generate the data. Dashed and dotted lines denote the 95-percentile width of a lognormal distribution with *σ* =0.1, and 0.5, respectively.

Most evolutionary rate estimates from the simulations with low evolutionary rate, condition (ii), had 95% HPD intervals that included the true mean value used to generate the data, 10^−5^ subs/site/unit time, at the expense of very wide 95% HPD intervals, compared with those in condition (i). Our analyses of data sets with a high evolutionary rate and narrow sampling window, condition (iii), had HPD intervals that were wider than those for condition (i), but narrower than those of condition (ii). All replicates with *σ*=0.0 or 0.1 had estimates that included the true mean value used to generate the data. In contrast, three data sets with *σ* =0.5 analysed under a strict clock yielded HPD intervals that did not include the true value. For data generated under *σ*=1.0, seven analyses under the strict clock and three under the relaxed clock also failed to recover the true value, although they always overlapped with the 95-percentile width of a lognormal distribution with mean 5×10^−3^ and *σ*=0.5. Analyses of the data with low evolutionary rate and narrow sampling window produced estimates that always included the true value of 10^−3^ subs/site/unit time in every case, but with very high uncertainty (fig. 4).

##### Comparison with Root-to-tip Regression

Using a subset of the heterochronous data sets, we conducted root-to-tip regression using phylogenetic trees inferred using maximum likelihood as implemented in PhyML 3.1 (Guindon et al. 2010) with the same substitution model as in our BEAST analyses, and with the placement of the root chosen to maximize *R*^2^ in TempEst (Rambaut et al. 2016). We selected data sets generated with a high evolutionary rate and with both constant and narrow sampling windows. Because GSS and NS correctly detected temporal signal under these conditions, these regressions demonstrate the extent to which this informal regression assessment matches the BETS approach. We did not attempt to provide a thorough benchmarking of the two methods here.

All regressions had *R*^2^ values that matched our expectation from the degree of among-lineage rate variation, that is, higher values of *σ* corresponded to lower values of *R*^2^ (fig. 5). The data with a wide sampling window yielded regression slopes ranging from 7.3×10^−3^ to 5.4×10^−3^ subs/site/unit time, which is similar to the evolutionary rate values used to generate the data. Although the root-to-tip regression is sometimes used to assess temporal signal, it has no cut-off values to make this decision. This becomes critical when considering the data with a narrow sampling window, for which the *R*^*2*^ was between 0.13 and 0.02. For example, the regression for a data set with *σ*=1 and narrow sampling window had an *R*^2^ of 0.02, which is sometimes considered sufficiently low as to preclude molecular clock analyses (Rieux and Balloux 2016). However, BETS supported temporal signal under a relaxed clock, with a log Bayes factor of 5.48 for this particular data set, which matches the simulation conditions. More importantly, even with such high rate variation, the evolutionary rate estimated using a relaxed clock and the correct sampling times included the true value used to generate the data (5×10^−3^ subs/site/unit time), with a 95% HPD interval of 2.15×10^−3^ to 1.90×10^−2^ subs/site/unit time, while the regression slope was 2.22×10^−2^ subs/site/unit time. A key implication of these comparisons is that BETS provides a formal assessment of temporal signal, unlike statistics computed from the regression. Moreover, the root-to-tip regression appears to be uninformative when the data have been sampled over a narrow time window and there is some rate variation among lineages.

**FIG. 5.**
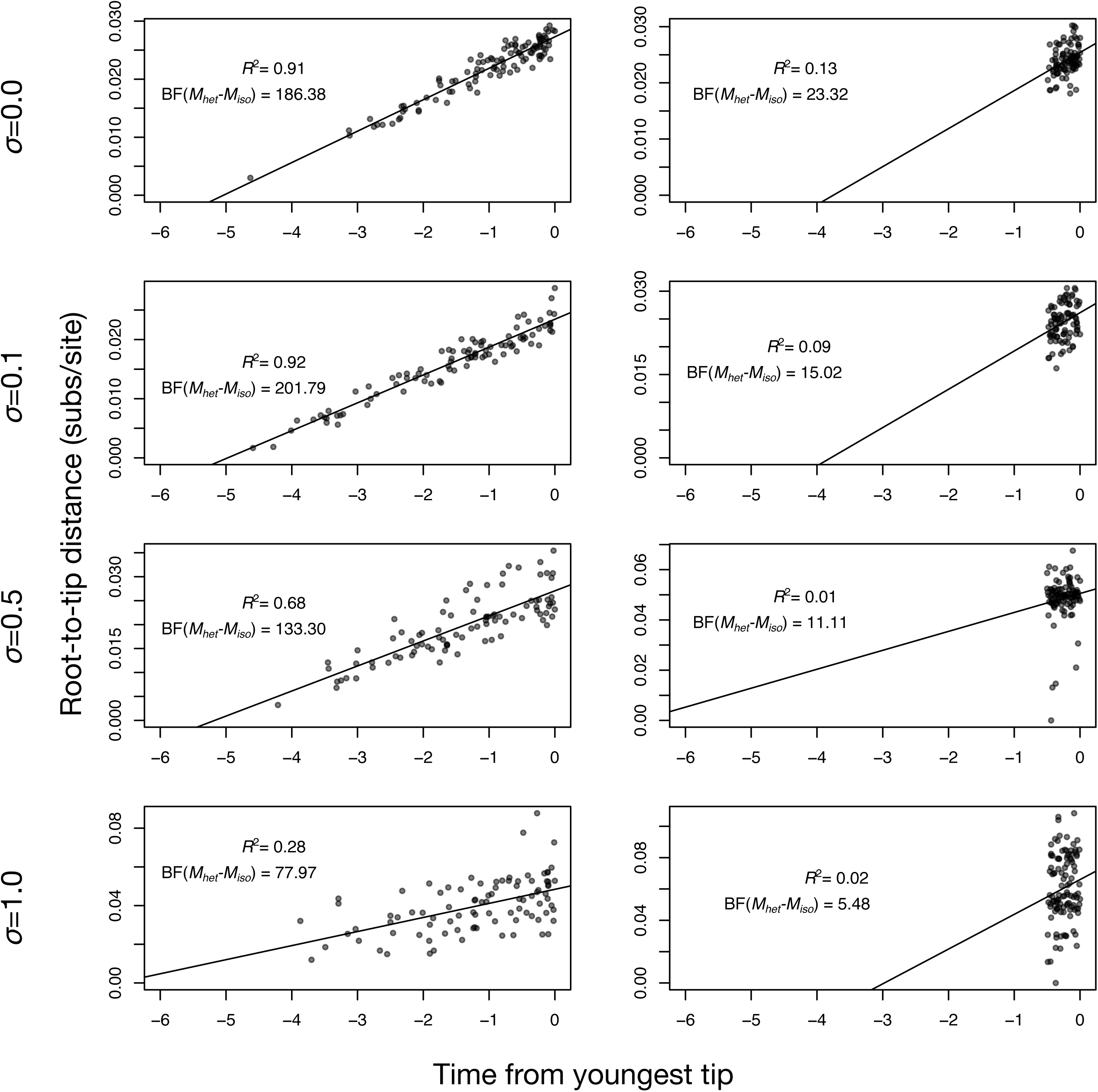
Root-to-tip regressions for a subset of data sets simulated with varying degrees of among-lineage rate variation (governed by the standard deviation *σ* of a lognormal distribution), using a high evolutionary rate and either a wide or narrow sampling window. The *y*-axis is the root-to-tip distance and the *x*-axis is the time from the youngest tip, where 0 is the present. Each point corresponds to a tip in the tree and the solid line is the best-fit linear regression using least-squares. The coefficient of determination, *R*^2^, is shown in each case. For comparison, the log Bayes factors of the best heterochronous model relative to the best isochronous model, BF(*M*_het_ -*M*_iso_), are also shown.

##### Simulations with phylo-temporal clustering

Phylo-temporal clustering sometimes occurs in empirical data due to limited opportunities for sample collection or varying degrees of population structure. We investigated the effects of phylo-temporal clustering by performing an additional set of simulations in which we specified five clades of 20 tips. To generate heterochronous data within each clade we set five possible sampling times that corresponded to the quantiles of sampling times from a birth-death process with the same exponential growth rate as in our birth-death simulations. We simulated trees conditioned on these clades and their sampling times. To generate the sequence data, we set*σ*=0.0 and *σ*=1.0. We estimated log marginal likelihoods using only GSS, owing to its accuracy.

Using GSS, BETS correctly identified temporal signal and the correct clock model in all simulations of heterochronous data. However, evolutionary rates were often overestimated for these data (fig. 6), a pattern that has been demonstrated previously (Duchêne et al. 2015; Murray et al. 2015). When the data were isochronous, BETS has lower performance, identifying the correct model in eight cases when *σ*=0.0 and seven cases when *σ*=1.0 (supplementary fig. S7, Supplementary Material online).

**FIG. 6.**
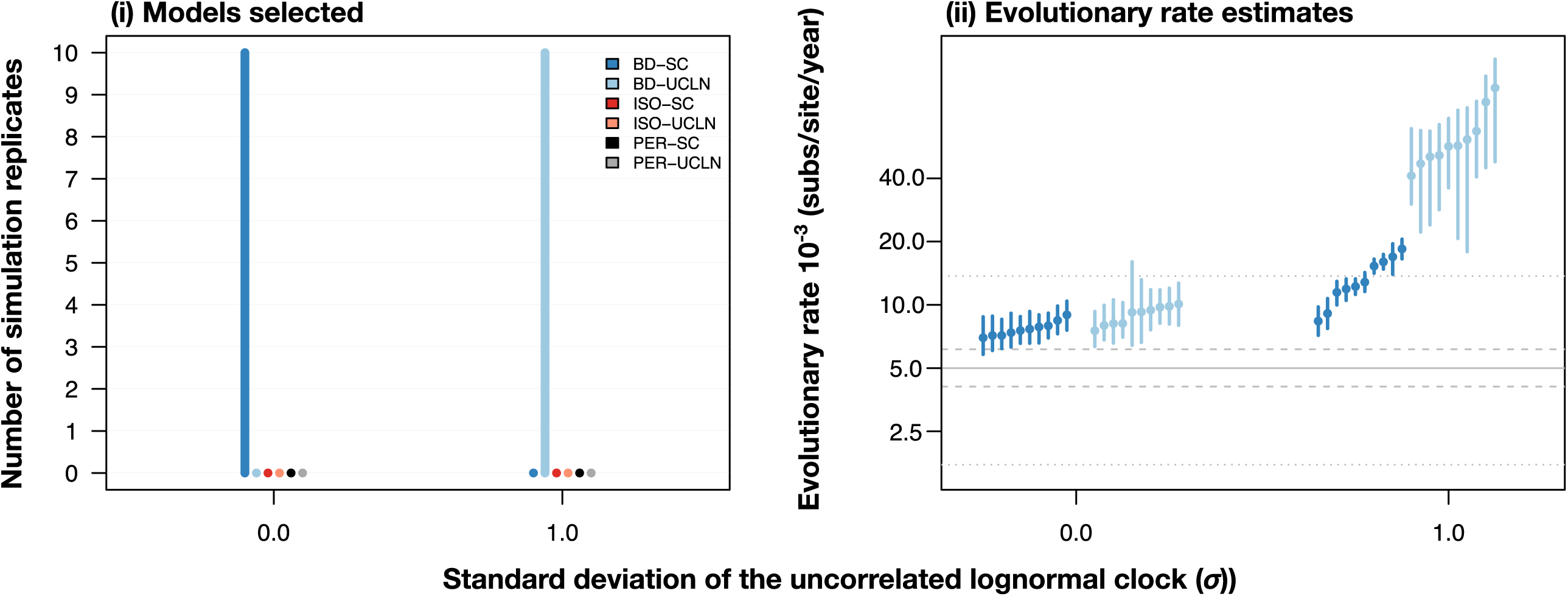
Results for heterochronous simulations with phylo-temporal clustering. The right-hand panel denotes models selected using generalized stepping-stone sampling under two degrees of among-lineage rate variation as determined by the standard deviation of a lognormal distribution,*σ* (along the *x*-axis). Each set of bars corresponds to a model and their height (along the *y*-axis) represents the number of times each model was selected out of ten simulation replicates. The bars are coloured depending on the analyses settings with two molecular clock models, strict clock (SC) and the uncorrelated relaxed clock with an underlying lognormal distribution (UCLN), and three settings for sampling times: generated under the birth-death process (BD), identical sampling times (Isochronous; ISO), and permuted (Permuted; PER). The left-hand panel shows evolutionary rate estimates for with correct sampling times using a strict clock and an uncorrelated relaxed clock with an underlying lognormal distribution.

##### Sensitivity and specificity

We investigated the extent to which detecting temporal signal could improve by using different cut-offs for the log Bayes factors. From a practical point of view, the main concern is that a data set with no temporal signal, for example when simulated here under isochronous trees, would be classified as heterochronous (i.e., false positives), resulting in spurious estimates of evolutionary rates and times. This problem was apparent in our simulations with a low evolutionary rate, where a number of isochronous data sets were classified as heterochronous. To determine such a possible cut-off value, we fit receiver operating characteristic (ROC) curves and calculated sensitivity and specificity (i.e., true positive and true negative rates, respectively).

Our simulations with high evolutionary rates were correctly classified, with sensitivity and specificity of 1.0 (fig. 7). Those with low evolutionary rates had a sensitivity and specificity of 0.68 and 0.85 with a wide sampling window and of 0.68 and 0.45 with a narrow sampling window. Importantly, these values correspond to a log Bayes factor cut-off optimized in the ROC curve fitting and is determined to be 1.04 for the simulations with a wide sampling window and 0.16 for those with a narrow sampling window. A more conservative approach to guard against false positives is to consider a higher cut-off value. A log Bayes factor of 3 is generally considered to be ‘strong’ evidence in favour of a model (Kass and Raftery 1995). In our simulations with low evolutionary rate this cut-off results in a specificity of 0.95, meaning that 95% of isochronous data sets were classified as such, at the expense of a low sensitivity of 0.43 for the data simulated with a wide sampling window, and of 0.0 for those with a narrow sampling window (note that sensitivity for the simulations with a low evolutionary rate and narrow sampling window using Bayes factor cut-off of 0.0 is already low, at 0.68). Importantly, using a log Bayes factor cut-off of 3 would still result in a specificity and sensitivity of 1.0 in our simulations with a high evolutionary rate.

**FIG. 7.**
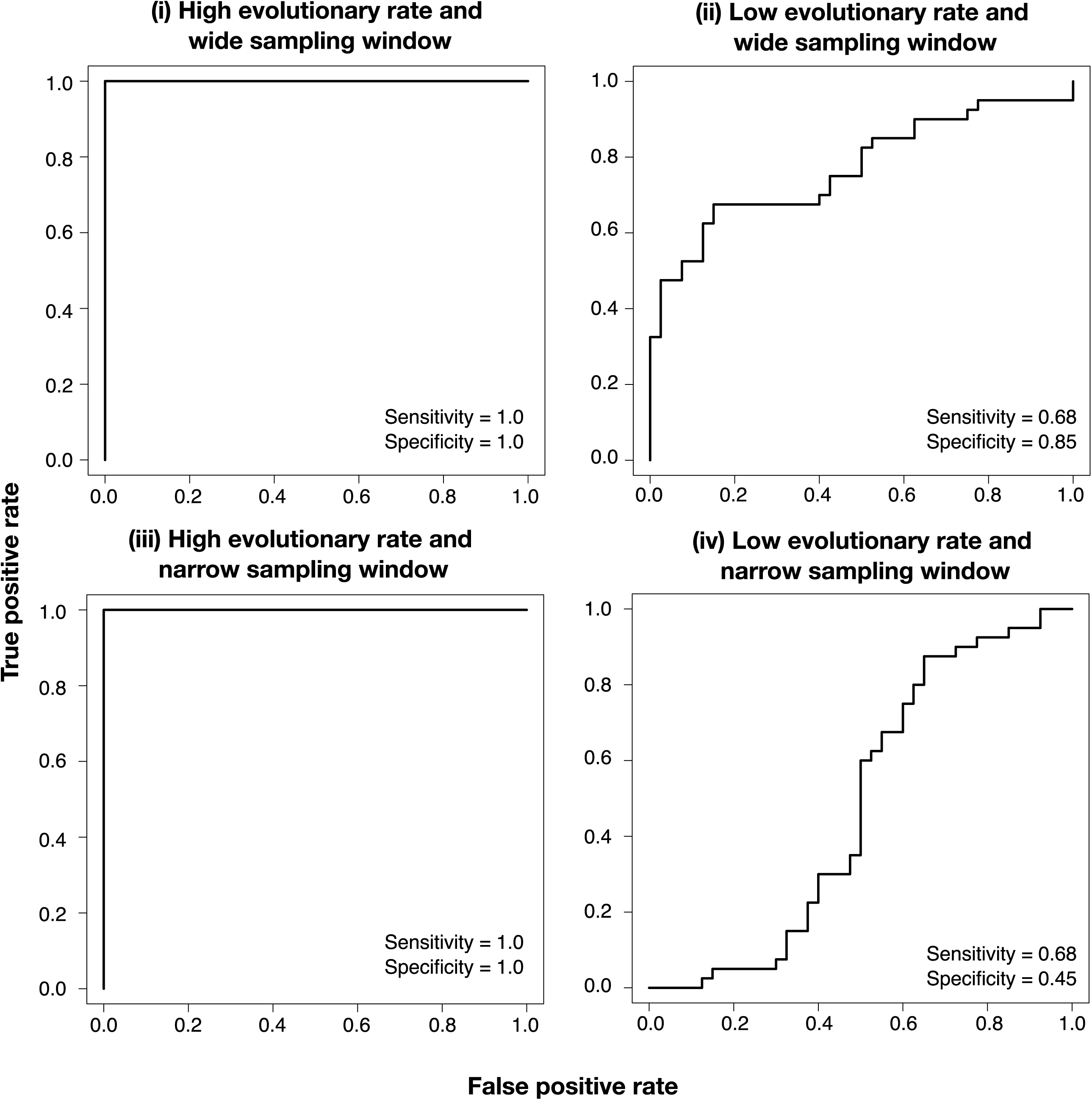
Receiver operating characteristic (ROC) curves for data simulated with high evolutionary rate and wide sampling window (i), low evolutionary rate and wide sampling window (ii), high evolutionary rate and narrow sampling window (iii), and low evolutionary rate and narrow sampling window (iv). Sensitivity and specificity values are shown in each case.

A key point about our data sets simulated with a low evolutionary rate is that they contain (very) low numbers of variable sites and unique site patterns (varying between 4 and 13), which can make model selection challenging. In order to increase accuracy, one could invest significant computational efforts to reduce estimator variance when repeated analyses prove inconclusive. The log Bayes factors for these data are much lower than for those generated using a higher evolutionary rate. We conducted another set of simulations with the same low evolutionary rate, but with much longer sequence alignments (10,000 nucleotides) to increase the number of variable sites and unique site patterns. For these longer alignments, the ROC curve indicated better performance of BETS, with sensitivity and specificity both equal to 0.83 with an optimal log Bayes factor of 1.39 (fig 8).

**FIG. 8.**
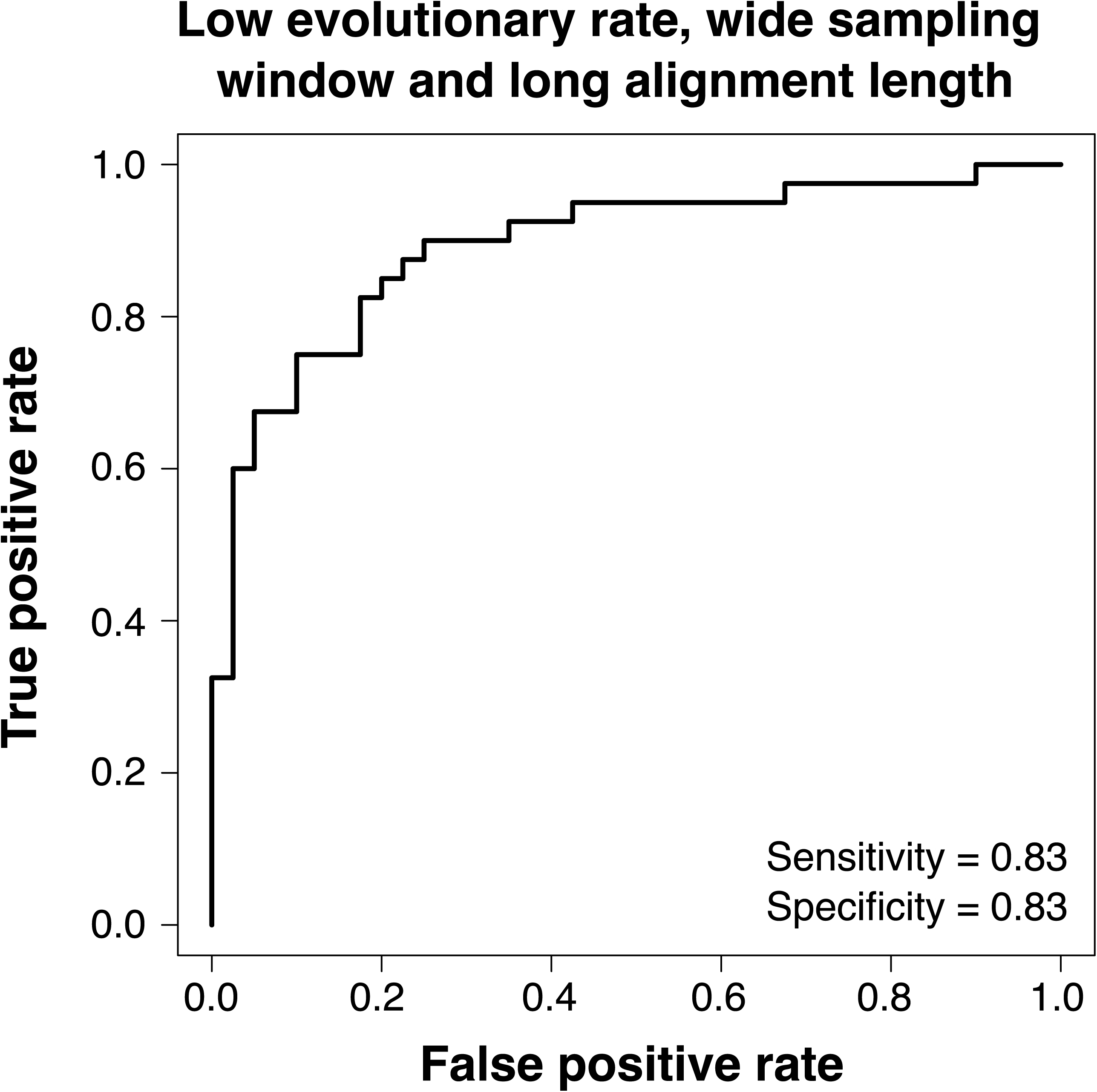
Receiver operating characteristic (ROC) curves for data simulated with low evolutionary rate, wide sampling window, and long sequence length (10,000 nucleotides). Sensitivity and specificity values are shown.

##### Analyses of Empirical Data Sets

We analysed five empirical data sets with similar configurations of sampling times as in our simulation study (Table 1). Two data sets consisted of rapidly evolving pathogens: *A/H1N1 influenza virus* (Hedge et al. 2013) and *Bordetella pertussis* (Bart et al. 2014). We also analysed a data set with highly divergent sequences of coronaviruses (Wertheim et al. 2013), and two data sets with ancient DNA: *Hepatitis B virus* (Patterson Ross et al. 2018) and mitochondrial genomes of dog species (Thalmann et al. 2013). Due to the demonstrated higher accuracy of GSS over NS (Fourment et al. 2019), we applied the BETS approach using the former method only.

**Table 1.** Details of empirical data sets used in this study.

| Data set | Number of sites<br>(nucleotides) | Number of samples | Sampling time range | Reference |
| --- | --- | --- | --- | --- |
| *A/H1N1 influenza virus* | 13,154 | 329 | 10 months (March to December 2009) | Hedge et al. (2013) |
| *Bordetella pertussis* | $4.9 \times 10^6$ | 150 | 89 years (1920 to 2009) | Bart et al. (2014) |
| Coronaviruses | 1,860 | 43 | 70 years (1941 to 2011) | Wertheim et al. (2013) |
| *Hepatitis B virus* | 3,271 | 137 | 445 years (2103 to 1568) | Patterson Ross et al. (2018) |
| Dog mtDNA | 14,596 | 50 | 36,000 years (to the present) | Thalmann et al. (2013) |

The *A/H1N1 influenza virus* data demonstrated clear temporal signal, with the strict clock and relaxed clock with the correct sampling times having the highest log marginal likelihoods, and a log Bayes factor of *M*_het_ with respect to *M*_iso_ of 150 (fig. 9). The strict clock had higher support than the relaxed clock for the correct sampling times (log Bayes factor 3.41). Broadly, this result is consistent with previous evidence of strong temporal signal and clocklike behaviour in this data set (Hedge et al. 2013). Using the strict clock with correct sampling times we estimated an evolutionary rate of 3.37×10^−3^ subs/site/year (95% HPD: 2.98×10^−3^ to 3.78×10^−3^).

**FIG. 9.**
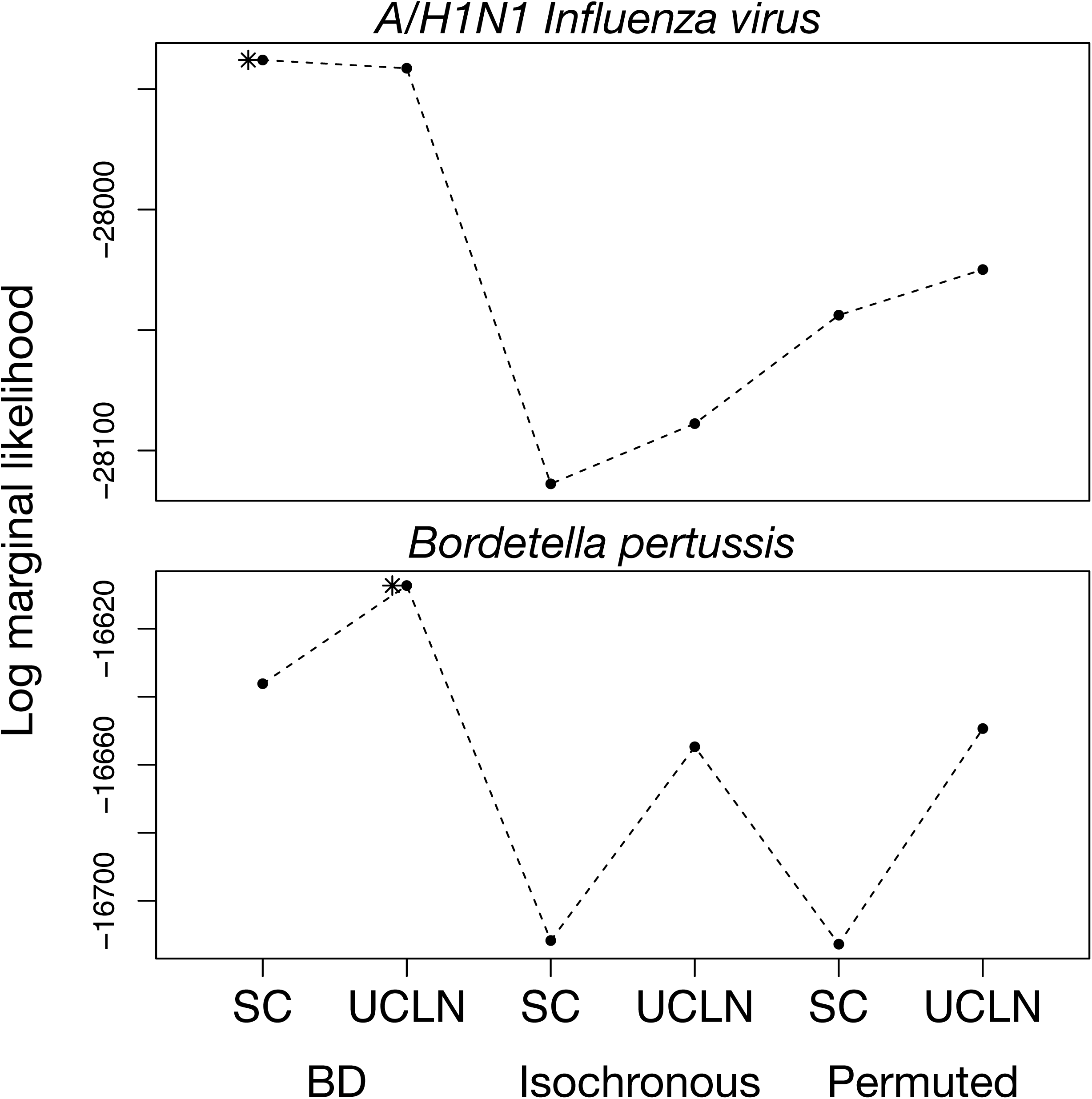
Log marginal likelihoods estimated using generalized stepping-stone sampling for six analysis settings for sequence data from rapidly evolving pathogens, *A/H1N1 Human influenza virus* and *Bordetella pertussis*. The *y*-axis is the log marginal likelihood and the *x*-axis shows the analysis settings, with two clock models, strict clock (SC) and the uncorrelated relaxed clock with an underlying lognormal distribution (UCLN), and three settings for sampling times: generated under the birth-death process (BD), identical sampling times (Isochronous), and permuted (Permuted). Solid points and dashed lines correspond to the log marginal likelihood estimates. The asterisk denotes the model with the highest log marginal likelihood.

We detected temporal signal in the *Bordetella pertussis* data set (fig. 9). The relaxed clock with the correct sampling times generated the highest log marginal likelihood, with a log Bayes factor relative to the strict clock of 28.86. The log Bayes factor for *M*_het_ relative to *M*_iso_ was 47.40. These results echo previous assessments of these data using a date-randomization test (Duchene et al. 2016). We estimated a mean evolutionary rate using the best model of 1.65×10^−7^ subs/site/year (95% HPD: 1.36×10^−7^ to 2.00×10^−7^).

Our analyses did not detect temporal signal in the coronavirus data, for which the strict clock and relaxed clock with no sampling times had the highest log marginal likelihoods. The log Bayes factor of *M*_het_ relative to *M*_iso_ was -16.82, indicating very strong support for the isochronous model. The relaxed clock was supported over the strict clock, with a log Bayes factor of 19.25 (fig. 10). The lack of temporal signal precludes any interpretation of our estimates of evolutionary rates and timescales. Previous analyses of these data suggested an ancient origin for this group of viruses using a substitution model that accounts for the effect of purifying selection over time (Wertheim et al. 2013), a model that we did not use here.

**FIG. 10.**
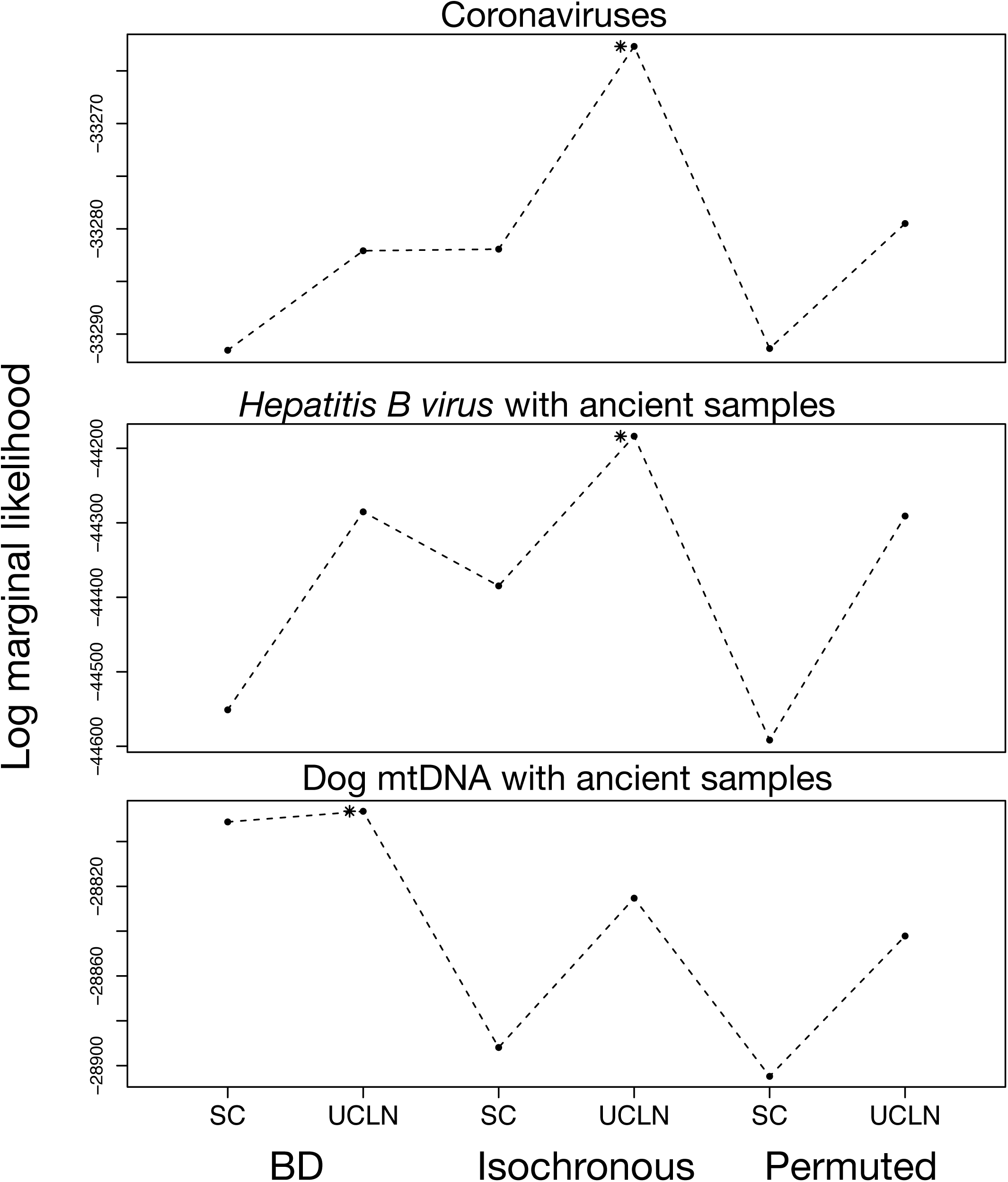
Log marginal likelihoods estimated using generalized stepping-stone sampling for six analysis settings for data sets with ancient DNA or highly divergent sequences. The *y*-axis is the log marginal likelihood and the *x*-axis shows the analysis settings, with two clock models, strict clock (SC) and the uncorrelated relaxed clock with an underlying lognormal distribution (UCLN), and three settings for sampling times: generated under the birth-death process (BD), identical sampling times (Isochronous), and permuted (Permuted). Solid points and dashed lines correspond to the log marginal likelihood estimates. The asterisk denotes the model with the highest log marginal likelihood.

The *Hepatitis B viru*s data set included several human genotypes with complete genomes, where 135 were modern sequences collected from 1963 to 2013 and two were ancient samples from human mummies from the 16^th^ century. Previous studies have not found any temporal signal in these data using different approaches, despite the inclusion of ancient sequences. Our estimates of log marginal likelihoods were consistent with a lack of temporal signal, with a log Bayes factor of -101.51 for *M*_het_ relative to *M*_iso_.

The dog mitochondrial genome data contained samples from up to 36,000 years before the present. BETS detected temporal signal in these data, with a log Bayes factor of 38.77 for *M*_het_ relative to *M*_iso_; this result is consistent with that of a date-randomization test in a previous study (Tong et al. 2018). The estimated evolutionary rate for these data using the best model had a mean of 1.08×10^−7^ subs/site/year (95% HPD: 7.49×10^−8^ to 1.52×10^−7^).

## Discussion

We have proposed BETS, a method that explicitly assesses the statistical support for including sequence sampling times in a Bayesian framework. It is a test of the presence of the temporal signal in a data set, which is an important prerequisite for obtaining reliable inferences in phylodynamic analyses. BETS considers the model ensemble, such that the method can detect temporal signal using models that account for evolutionary rate variation among lineages. The results of our analyses demonstrate that our method is effective in a wide range of conditions, including when the evolutionary rate is low or when the sampling window represents a small portion of the timespan of the tree.

BETS does not require date permutations, which sets it apart from the widely used date-randomization test for temporal structure. Date-randomization tests address the question of whether a particular association between sequences and sampling times produces estimates different from those obtained from data sets with permuted sampling times (Duchêne et al. 2015; Murray et al. 2015). However, such an approach is not a formal test of temporal signal in the data because the permutations do not necessarily constitute an appropriate null model. Because our method does not require permutations, it has the benefit of being robust to using a limited number of sampling times.

Accurate calculations of log marginal likelihoods are essential for BETS. In our simulation study, we found that GSS and NS correctly assessed the presence and absence of temporal signal in the data under most conditions. The correct clock model was also identified, although in a few instances NS preferred an overparameterized model. Conceivably, using different log marginal likelihood estimators might affect the actual model selected. Murray et al. (2015) also employed a Bayesian model-testing approach using the AICM to assess temporal signal. In their study, the AICM performed well in simulations, but failed to detect temporal signal in empirical data. We attribute this finding to the low accuracy of AICM compared with path sampling methods (Baele et al. 2012, 2013), and suggest careful consideration of the log marginal likelihood estimator for tests of temporal signal. In a recent review, Fourment et al. (2020) found GSS to be a highly accurate albeit computationally demanding log marginal likelihood estimator.

A key benefit of BETS is that the complete model is considered. It is straightforward to use any model for which the log marginal likelihood can be calculated, including other models of among-lineage rate variation, unlike in simpler data exploration methods such as root-to-tip regression. In the particular case of local clock models (Drummond and Suchard 2010; Worobey et al. 2014; Bletsa et al. 2019), the root-to-tip regression is uninformative because it assumes that the slope represents a single mean evolutionary rate.

We find that highly precise and accurate evolutionary rate estimates are associated with strong Bayes factor support for heterochronous models (fig. 4 and supplementary fig. 3, Supplementary Material Online). Bayes factors provide a coherent approach to identifying the presence of temporal signal, instead of providing a potentially subjective gradient of strength of such signal. In contrast, root-to-tip regression offers an important visual aid for uncovering problems with data quality and to inspect clocklike behaviour, but the absence of appropriate statistics means that there is no clear objective way of determining whether the data contain temporal information. Consider the regressions in figure 5 for data with a high evolutionary rate and narrow sampling window. Even when among-lineage rate variation is low (σ=0.1), the data points form a cloud, with a low R^2^ of 0.09. However, the apparent ‘noise’ around the regression line is probably the result of stochasticity in sequence evolution and of the narrow sampling window relative to the age of the root of the tree. In fact, for this particular data set the model with the highest log marginal likelihood is the strict clock with correct sampling times.

In all of our analyses, we ensured that the priors for different models and configurations of sampling times were identical because, as with all Bayesian analyses, model comparison using log marginal likelihoods can depend on the choice of prior (Oaks et al. 2019). For example, the tree prior can affect inferences of temporal signal, as it is part of the full model specification. Here we used an exponential-growth coalescent tree prior, which closely matches the demographic dynamics of the birth-death process under which the data were simulated. The effect of using an inappropriate tree prior on tests of temporal signal requires further investigation, but previous studies have suggested that there is only a small impact on estimates of rates and times if the sequence data are informative (Ritchie et al. 2017; Möller et al. 2018).

An interesting finding is that statistical support for isochronous sampling times in truly isochronous data is lower than that for the correct sampling times in truly heterochronous data. This can potentially lead to an increased risk of incorrectly concluding the presence of temporal signal. In particular, in isochronous data simulated with a low evolutionary rate, and with very few variable sites, the best models were sometimes those that included sampling times, albeit with very low log Bayes factors (e.g., supplementary fig. S1 and fig. S2, Supplementary Material online). This probably occurs because stochastic error associated with a small amount of evolution leads to low power for model selection. While increasing the computational settings for (log) marginal likelihood estimation can alleviate these issues, this may not be feasible when analysing large data sets. Further, our sensitivity and specificity analyses demonstrate that a practical way to address this problem is to use a more conservative log Bayes factor cut-off of 3 as evidence of temporal structure, as opposed to simply choosing the model with the highest marginal likelihood. This cut-off matches ‘strong’ evidence in favour of a model as suggested by Kass and Raftery (1995).

Permuting sampling times led to poor model fit, as expected. This procedure has substantial computing requirements, depending on the number of permutations that are performed, and we find that such date permutations are of limited value for model testing when the data are highly informative (e.g., figs. 2 and 3). However, in data sets with very low information content, such as those that were produced by simulations with a low evolutionary rate here, conducting a small number of date permutations might offer a conservative approach to determining whether model fit and parameter estimates are driven by a particular set of sampling times, as one would expect in the presence of temporal signal.

The nature of the BETS approach means that every parameter in the model has a prior probability, including the evolutionary rate. Because evolutionary rates and times are nonidentifiable, it is conceivable that an informative prior on the rate or on the age of an internal node might have a stronger effect than the sampling times on the posterior, for example if the samples span a very short window of time. Such analyses with informative evolutionary rate priors effectively include several simultaneous sources of calibration information (i.e., sampling times, internal nodes, and an informative rate prior). Using sampling times in addition to other sources of calibration information might still be warranted if such external sources of information are available.

Most of our heterochronous simulations yielded evolutionary rate estimates that contained the true value used to generate the data, indicative of the accuracy of our estimations. However, it is important to note that all tests of temporal signal, including BETS, aim to determine whether there is an association between genetic divergence and time, which is not equivalent to asking whether evolutionary rate estimates are accurate, a question that depends on information content of the data and the extent to which the model describes the process that generated the data. Phylo-temporal clustering is a particular situation where temporal information in the data is very limited, leading to an upward bias in the evolutionary rate (Murray et al. 2015), even in the presence of temporal signal. As such, investigating the degree of phylo-temporal clustering is an important step prior to interpreting any inferences made using the molecular clock (Duchêne et al. 2016; Tong et al. 2018).

Analyses with multiple calibrations can also allow uncertainty in sequence sampling times, especially in data sets that include ancient DNA, where sampling times can be treated as parameters in the model (Shapiro et al. 2011). BETS provides a coherent approach for assessing temporal structure in these circumstances, unlike date-randomization tests that typically use point values for sampling times. In fact, BETS can be used as a means to validate whether a sample is modern or ancient.

In general, the increasing adoption of Bayesian model testing in phylogenetics has great potential for improving our confidence in estimates of evolutionary rates and timescales. The test that we have proposed here, BETS, provides a coherent and intuitive framework to test for temporal information in the data.

## Materials and Methods

### Simulations

We simulated phylogenetic trees under a stochastic birth-death process using MASTER v6.1 (Vaughan and Drummond 2013), by specifying birth rate *λ*=1.5, death rate *μ*=0.5, and sampling rate *ψ*=0.5. This corresponds to an exponentially growing infectious outbreak with reproductive number *R*_0_=1.5 and a wide sampling window. We set the simulation time to 5 units of time, which corresponds to the time of origin of the process. For isochronous trees, we used similar settings, but instead of using the sampling rate, we sampled each tip with probability*ρ*=0.5 when the process was stopped after 5 units of time (i.e. *μ*=1.0 and *ψ*=0.0). Some of our analyses consisted of artificially specifying sampling times for isochronous trees, which we set to those that we would have obtained from a birth-death process with *μ*=0.5 and *ψ*=0.5.

In a second set of simulations of heterochronous trees, we generated trees with a narrow sampling window. We specified two intervals for *μ* and*ψ*. The first interval spanned 4.5 units of time with *μ*=1.0 and *ψ*=0.0, and the second interval 0.5 units of time with *μ*=0.1 and *ψ*=0.9. As a result, the process still had a constant become-uninfectious rate (*μ*+*ψ*), but samples were only collected in the second interval. The high sampling rate in the second interval resulted in trees with similar numbers of tips to those with a wide sampling window, but where their ages only spanned 0.5 units of time.

We only considered the simulated trees that contained between 90 and 110 tips. The trees generated in MASTER are chronograms (with branch lengths in units of time), so we simulated evolutionary rates to generate phylograms (with branch lengths in units of subs/site). To do this we specified the uncorrelated lognormal relaxed clock with a mean rate of 5×10^−3^ or 10^−5^ subs/site/unit time and a standard deviation *σ* of 0.0 (corresponding to a strict clock), 0.1, 0.5, or 1.0. We simulated sequence evolution along these phylograms under the HKY nucleotide substitution model (Hasegawa et al. 1985). We added among-site rate variation using a discretized gamma distribution (Yang 1994, 1996) using Phangorn v2.5 (Schliep 2011) to generate sequence alignments of 4,000 and 10,000 nucleotides. We set the transition-to-transversion ratio of the HKY model to 10 and the shape of the gamma distribution to 1, which is similar to estimates of these parameters in influenza viruses (Duchene et al. 2014; Hedge and Wilson 2014). For each simulation scenario we generated 10 sequence alignments.

To simulate data under phylo-temporal clustering we specified five clades with 20 tips each to generate trees of 100 tips. For the heterochronous data, we specified one of five possible sampling times for each clade, which corresponded to quantiles from a birth-death process as used in our simulations above. For the isochronous data we constrained the tips to have identical sampling times. We specified these clades and sampling times in BEAST as monophyletic groups and sampled trees from the prior under a coalescent process with exponential growth parameterized with *λ*=1.5 and *δ*=1, such that it has the same growth rate as the birth-death trees. We conducted these simulations under the coalescent, rather than the birth-death, because this process is typically conditioned on the number and age of samples, whereas the birth-death explicitly models sampling over time. We simulated sequence data sets as above, but in this case we only considered an evolutionary rate of 5×10^−3^ subs/site/year and a*σ* of 0.0 or 1.0.

### Estimation of Log Marginal Likelihoods Using Nested Sampling

We analysed the data in BEAST 2.5 using the matching substitution model, the exponential-growth coalescent tree prior, the strict clock or relaxed clock, and different configurations of sampling times. We chose the exponential-growth coalescent tree prior, instead of the birth-death tree prior, because it is conditioned on the samples instead of assuming a sampling process; this ensures that the marginal likelihoods for isochronous and heterochronous trees are comparable.

We specified proper priors on all parameters, which is essential for accurate estimation of log marginal likelihoods (Baele et al., 2013). In our heterochronous analyses the prior on the evolutionary rate had a uniform distribution bounded between 0 and 1. We made this arbitrary choice to set a somewhat uninformative prior and because the default prior in BEAST 2.5 is a uniform distribution between 0 and infinity, which is improper. Owing to the non-identifiability of evolutionary rates and times, neither can be inferred in the absence of calibrating information, so in our isochronous analyses we fixed the value of the evolutionary rate to 1. The initial NS chain length was chosen so as to draw 20,000 samples, with 20,000 steps, 32 particles, and a subchain length of 5,000 (note that NS is not equivalent to standard MCMC, nor is the definition of an iteration/step). The chain length and its accompanying sampling frequency were adjusted to obtain effective sample sizes for key parameters of at least 200 (computed in the NS output in BEAST 2.5). Examples of MASTER files and BEAST 2.5 input files for NS are available online (supplementary data, Supplementary Material online).

### Estimation of Log Marginal Likelihoods Using Generalized Stepping-Stone Sampling

We used BEAST 1.10 with the same model specifications and priors as in BEAST2, except for the prior on the evolutionary rate, for which we used the approximate continuous-time Markov chain (CTMC) reference prior (Ferreira and Suchard 2008). Because our simulation analyses of GSS and NS differ in this prior, the log marginal likelihood estimates are not directly comparable, so for each simulation we report log Bayes factors of competing models instead of the individual log marginal likelihoods. The GSS implementation in BEAST 1.10 has two different working priors for the tree generative process: a matching tree prior and a product of exponentials. The latter approach is the most generally applicable and is the one that we used here (Baele et al. 2016).

We used an initial MCMC chain length of 5×10^7^ steps sampling every 5000 steps. After discarding 10% of the samples obtained, the remaining samples were used to construct the working distributions for the GSS analysis through kernel density estimation. The log marginal likelihood estimation comprised 100 path steps distributed according to quantiles from a *β* distribution with *α*=0.3, with each of the 101 resulting power posterior inferences running for 5×10^5^ iterations. We assessed sufficient sampling for the initial MCMC analysis by verifying that the effective sample sizes for key parameters were at least 200 in Coda v0.19 (Plummer et al. 2006). If this condition was not met, we doubled the length of the MCMC and reduced sampling frequency accordingly. Examples of MASTER files and BEAST 1.10 input files for GSS are available online (supplementary data, Supplementary Material online).

### Receiver Operating Characteristic (ROC) Curves

ROC curves are generated by plotting the true positive rate (TPR, i.e. the sensitivity) against the false positive rate (FPR, i.e. 1 – specificity) at a range of selected thresholds and allows assessment of the performance of a binary classifier system. We fit ROC curves to the different simulation scenarios using the R package ROCR (Sing et al. 2005). We classified data as ‘positives’ and ‘negatives’ if they were generated under a heterochronous or isochronous (i.e., no temporal signal) model, respectively. In order to determine the optimal cut-off value, we determined the point on the ROC curve closest to a TPR of 1 and an FPR of 0 (i.e. we assigned equal importance to sensitivity and specificity). We did not explore assigning different costs to false positives and false negatives.

### Analyses of Empirical Data Sets

We downloaded sequence alignments from their original publications (Table 1): complete genomes of the 2009 pandemic lineage of *A/H1N1 influenza virus* (Hedge et al. 2013), whole genome sequences of *B. pertussis* (Bart et al. 2014; Duchene et al. 2016), RdRP sequences of coronaviruses (Wertheim et al. 2013), complete genomes of *Hepatitis B virus* (Patterson Ross et al. 2018), and dog mitochondrial genomes (Thalmann et al. 2013). The data and BEAST input files are available in the Supplementary Material online.

Briefly, we used similar settings as in our simulations to estimate log marginal likelihoods using GSS. For sequence sampling times we considered the correct sampling times, no sampling times (i.e., isochronous), and permuted sampling times. We also specified tree priors as follows: an exponential-growth coalescent for the *A/H1N2 influenza virus, Bordetella pertussis*, coronaviruses, and *Hepatitis B virus* data sets, and a constant-size coalescent for the dog mitochondrial genomes as used by Tong et al. (2018). We again chose the HKY+Γ substitution model, except in the analysis of *Hepatitis B virus* data, for which we used the GTR+Γ model (Tavaré 1986), and in the analysis of the dog data set for which we used the SRD06 substitution model (Shapiro et al. 2006) for coding regions and the GTR+Γ for noncoding regions.

## Supplementary Material

Supplementary data are available online.

## Acknowledgements

We thank the Editor and two anonymous reviewers for useful comments on previous versions of this manuscript.

## Supplementary Material

**FIG. S1**. Models selected for isochronous data using generalized stepping-stone sampling under two evolutionary rates, shown in each panel and noted in the main text as conditions (i) and (ii), and four degrees of among-lineage rate variation as determined by the standard deviation of a lognormal distribution,*σ* (along the *x*-axis). Each set of bars corresponds to a model and their height (along the *y*-axis) represents the number of times each model was selected out of ten simulation replicates. The bars are colored depending on the analyses settings with two molecular clock models, strict clock (SC) and the uncorrelated relaxed clock with an underlying lognormal distribution (UCLN), and three settings for sampling times: generated under the birth-death process the using five quantiles (BD; i.e. correct sampling times with phylo-temporal clustering), identical sampling times (Isochronous; ISO), and permuted (Permuted; PER).

**FIG. S2**. Models selected for isochronous data using nested sampling under two evolutionary rates, shown in each panel and noted in the main text as conditions (i) and (ii), and four degrees of among-lineage rate variation as determined by the standard deviation of a lognormal distribution,*σ* (along the *x*-axis). Each set of bars corresponds to a model and their height (along the *y*-axis) represents the number of times each model was selected out of ten simulation replicates. The bars are colored depending on the analyses settings with two molecular clock models, strict clock (SC) and the uncorrelated relaxed clock with an underlying lognormal distribution (UCLN), and three settings for sampling times: generated under the birth-death process (BD), identical sampling times (Isochronous; ISO), and permuted (Permuted; PER).

**FIG. S3**. Log Bayes factors of heterochronous data simulated with a high evolutionary rate and a wide sampling window. Each panel shows the results for data sets simulated with a different degree of among-lineage rate variation, governed by the standard deviation *σ* of a lognormal distribution. The *x*-axis depicts six analysis settings, with two molecular clock models, strict clock (SC) and the uncorrelated relaxed clock with an underlying lognormal distribution (UCLN), and three settings for sampling times: generated under the birth-death process (BD), identical sampling times (Isochronous), and permuted (Permuted). The points have been jittered to facilitate visualization. The *y*-axis shows log Bayes factors relative to the best model. Black circles correspond to estimates using generalized stepping-stone sampling and grey circles correspond to estimates using nested sampling. We conducted 10 simulation replicates, with each replicate data set analysed under the six analysis settings and two marginal likelihood estimators, such that stochastic error might cause differences in the preferred model. The number next to each cloud of points denotes the number of times (out of 10) that the corresponding model had the highest log marginal likelihood with generalized stepping-stone sampling (in black) and nested sampling (in grey).

**FIG. S4**. Log Bayes factors of heterochronous data simulated with a low evolutionary rate and a wide sampling window. Each panel shows the results for data sets simulated with a different degree of among-lineage rate variation, governed by the standard deviation *σ* of a lognormal distribution. The *x*-axis depicts six analysis settings, with two molecular clock models, strict clock (SC) and the uncorrelated relaxed clock with an underlying lognormal distribution (UCLN), and three settings for sampling times: generated under the birth-death process (BD), identical sampling times (Isochronous), and permuted (Permuted). The points have been jittered to facilitate visualization. The *y*-axis shows log Bayes factors relative to the best model. Black circles correspond to estimates using generalized stepping-stone sampling and grey circles correspond to estimates using nested sampling. We conducted 10 simulation replicates, with each replicate data set analysed under the six analysis settings and two marginal likelihood estimators, such that stochastic error might cause differences in the preferred model. The number next to each cloud of points denotes the number of times (out of 10) that the corresponding model had the highest log marginal likelihood with generalized stepping-stone sampling (in black) and nested sampling (in grey).

**FIG. S5**. Log Bayes factors of heterochronous data simulated with a high evolutionary rate and a narrow sampling window. Each panel shows the results for data sets simulated with a different degree of among-lineage rate variation, governed by the standard deviation *σ* of a lognormal distribution. The *x*-axis depicts six analysis settings, with two molecular clock models, strict clock (SC) and the uncorrelated relaxed clock with an underlying lognormal distribution (UCLN), and three settings for sampling times: generated under the birth-death process (BD), identical sampling times (Isochronous), and permuted (Permuted). The points have been jittered to facilitate visualization. The *y*-axis shows log Bayes factors relative to the best model. Black circles correspond to estimates using generalized stepping-stone sampling and grey circles correspond to estimates using nested sampling. We conducted 10 simulation replicates, with each replicate data set analysed under the six analysis settings and two marginal likelihood estimators, such that stochastic error might cause differences in the preferred model. The number next to each cloud of points denotes the number of times (out of 10) that the corresponding model had the highest log marginal likelihood with generalized stepping-stone sampling (in black) and nested sampling (in grey).

**FIG. S6**. Log Bayes factors of heterochronous data simulated with a low evolutionary rate and a narrow sampling window. Each panel shows the results for data sets simulated with a different degree of among-lineage rate variation, governed by the standard deviation *σ* of a lognormal distribution. The *x*-axis depicts six analysis settings, with two molecular clock models, strict clock (SC) and the uncorrelated relaxed clock with an underlying lognormal distribution (UCLN), and three settings for sampling times: generated under the birth-death process (BD), identical sampling times (Isochronous), and permuted (Permuted). The points have been jittered to facilitate visualization. The *y*-axis shows log Bayes factors relative to the best model. Black circles correspond to estimates using generalized stepping-stone sampling and grey circles correspond to estimates using nested sampling. We conducted 10 simulation replicates, with each replicate data set analysed under the six analysis settings and two marginal likelihood estimators, such that stochastic error might cause differences in the preferred model. The number next to each cloud of points denotes the number of times (out of 10) that the corresponding model had the highest log marginal likelihood with generalized stepping-stone sampling (in black) and nested sampling (in grey).

**FIG S7**. Results for isochronous simulations with phylo-temporal clustering using generalized stepping-stone sampling under two degrees of among-lineage rate variation as determined by the standard deviation of a lognormal distribution,*σ* (along the *x*-axis). Each set of bars corresponds to a model and their height (along the *y*-axis) represents the number of times each model was selected out of ten simulation replicates. The bars are colored depending on the analyses settings with two molecular clock models, strict clock (SC) and the uncorrelated relaxed clock with an underlying lognormal distribution (UCLN), and three settings for sampling times: generated the birth-death process using five quantiles (BD; i.e. artificially producing phylo-temporal clustering), identical sampling times (Isochronous; ISO), and permuted (Permuted; PER).

