## Supplementary figures and images for "Bayesian Evaluation of Temporal Signal in Measurably Evolving Populations"

### Supplementary material

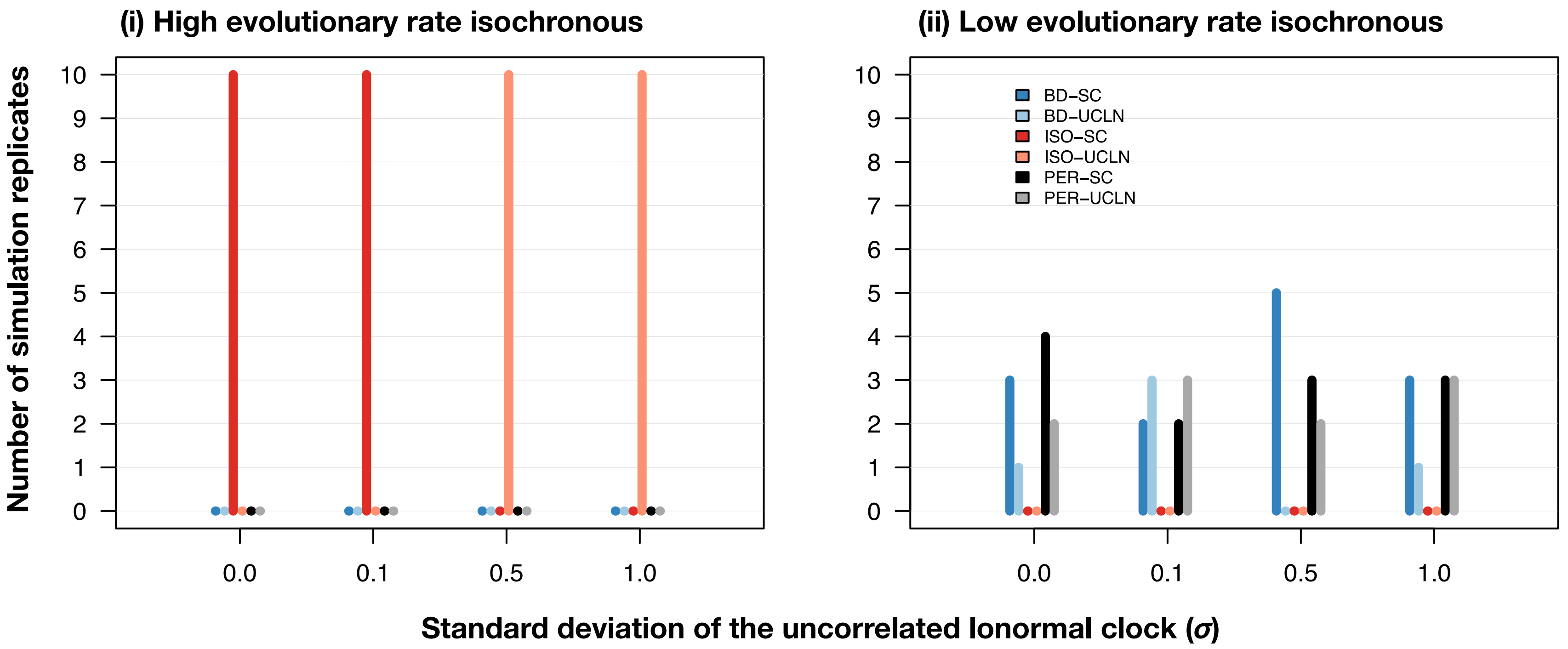

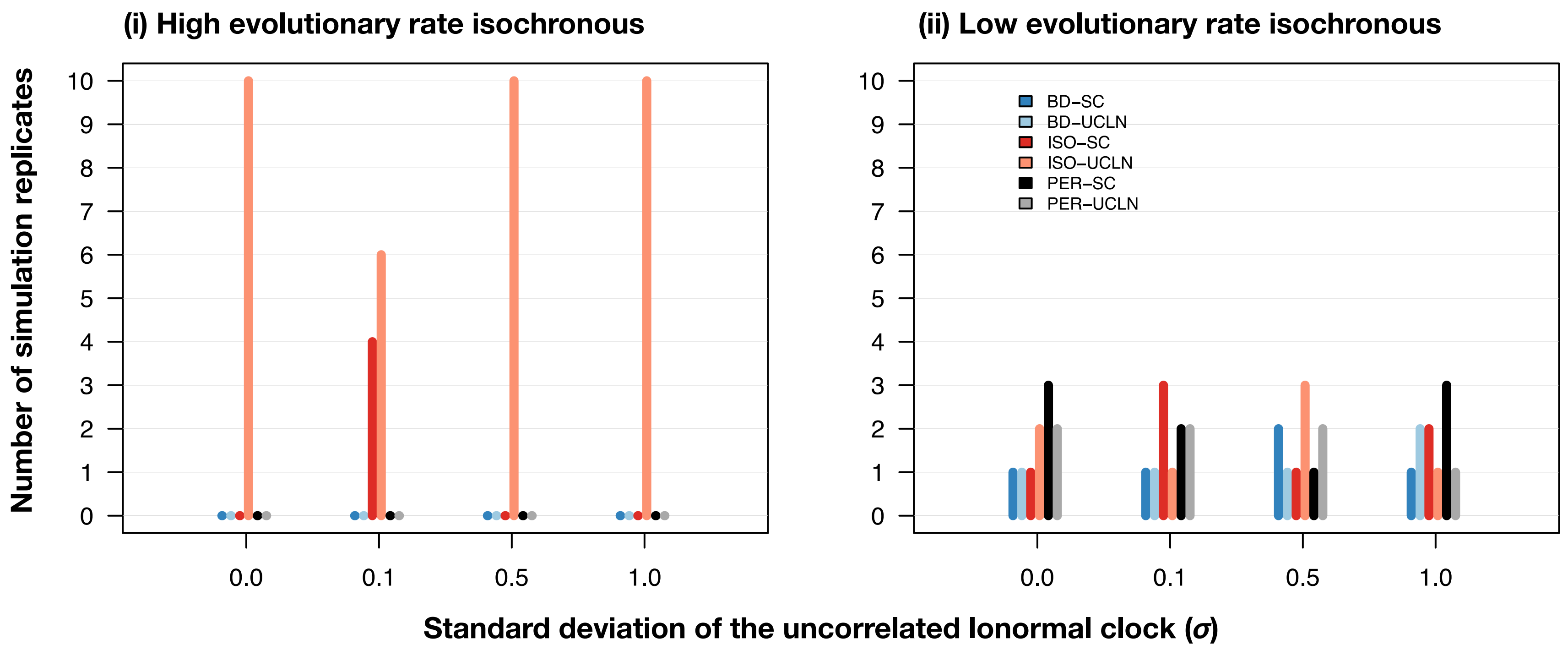

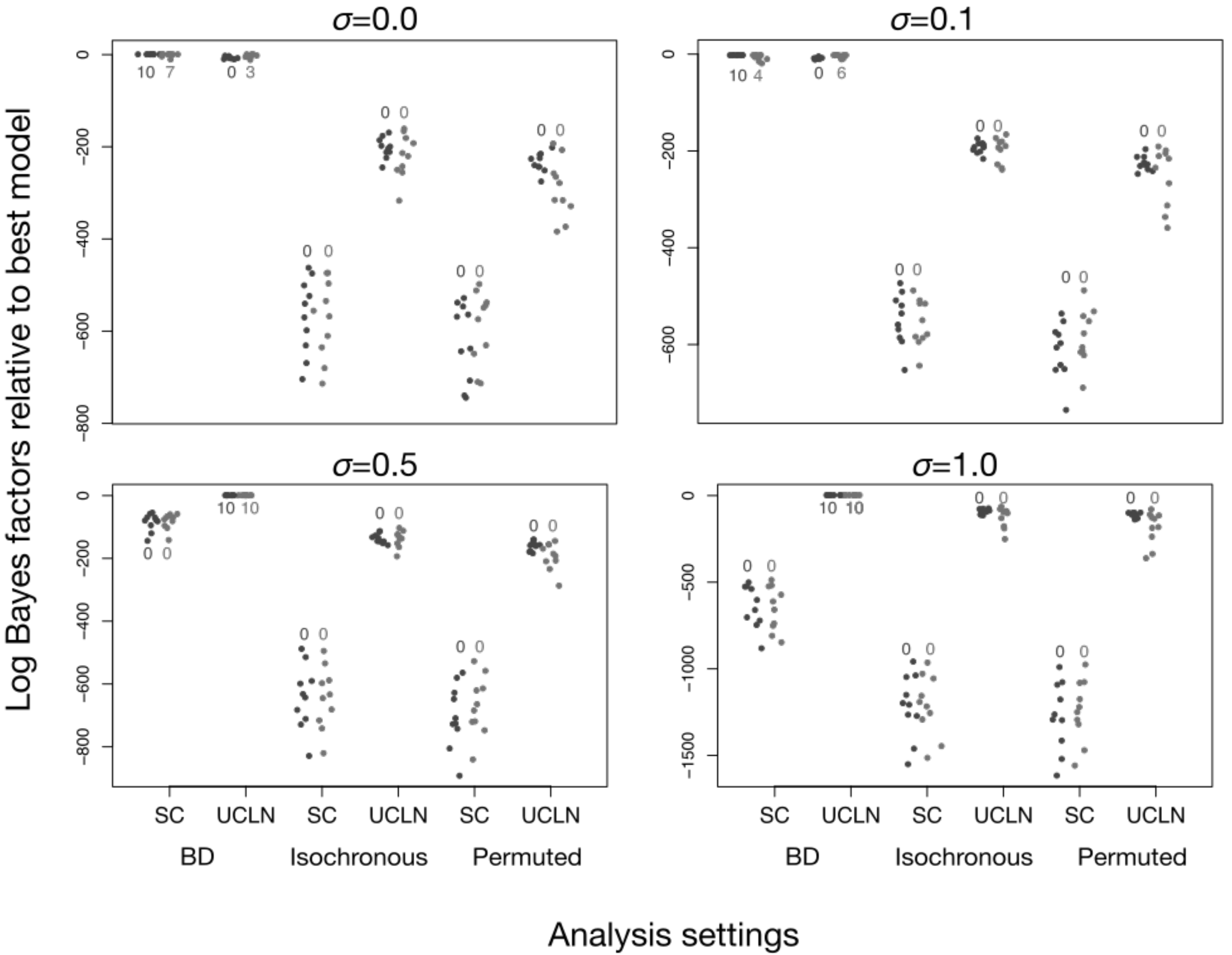

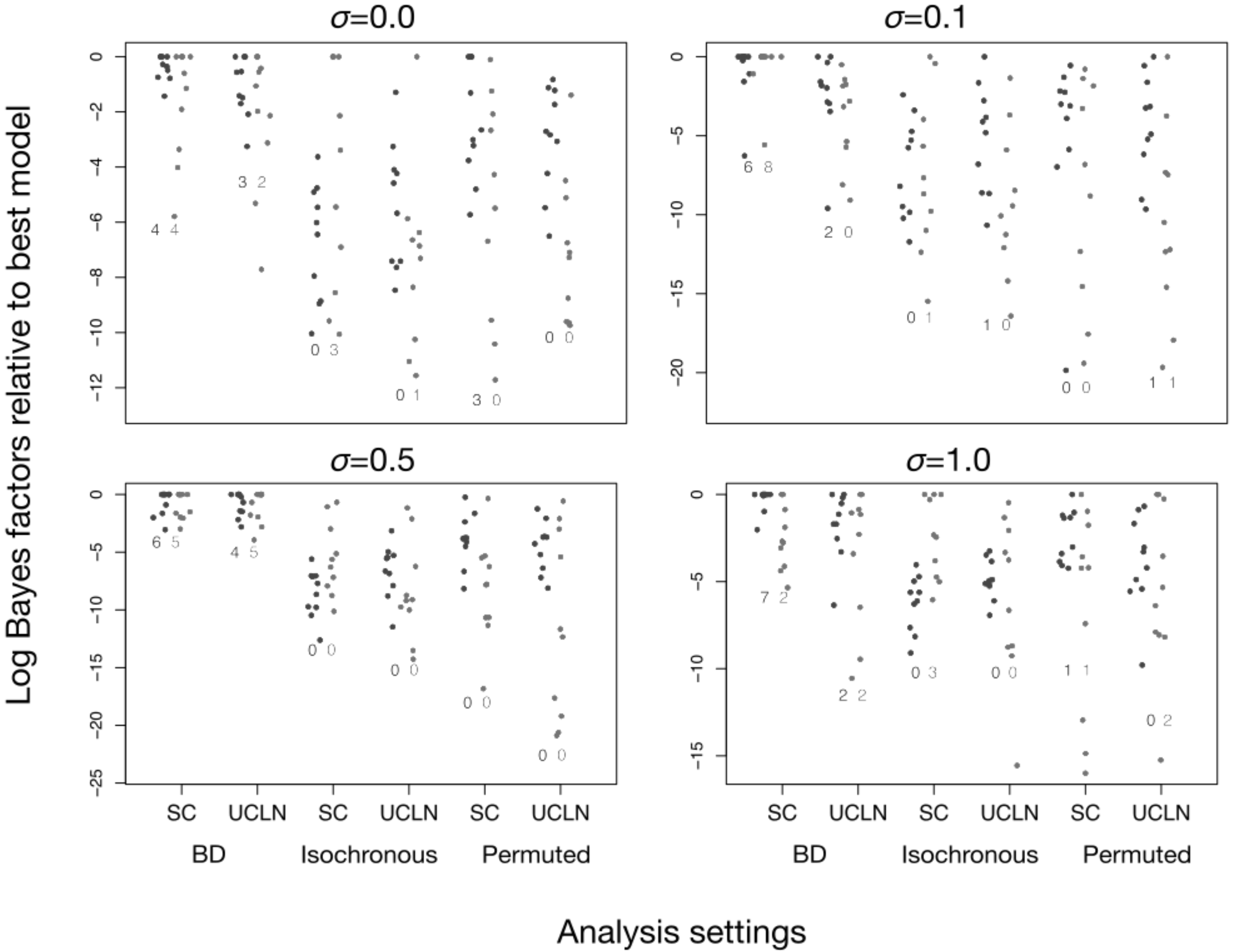

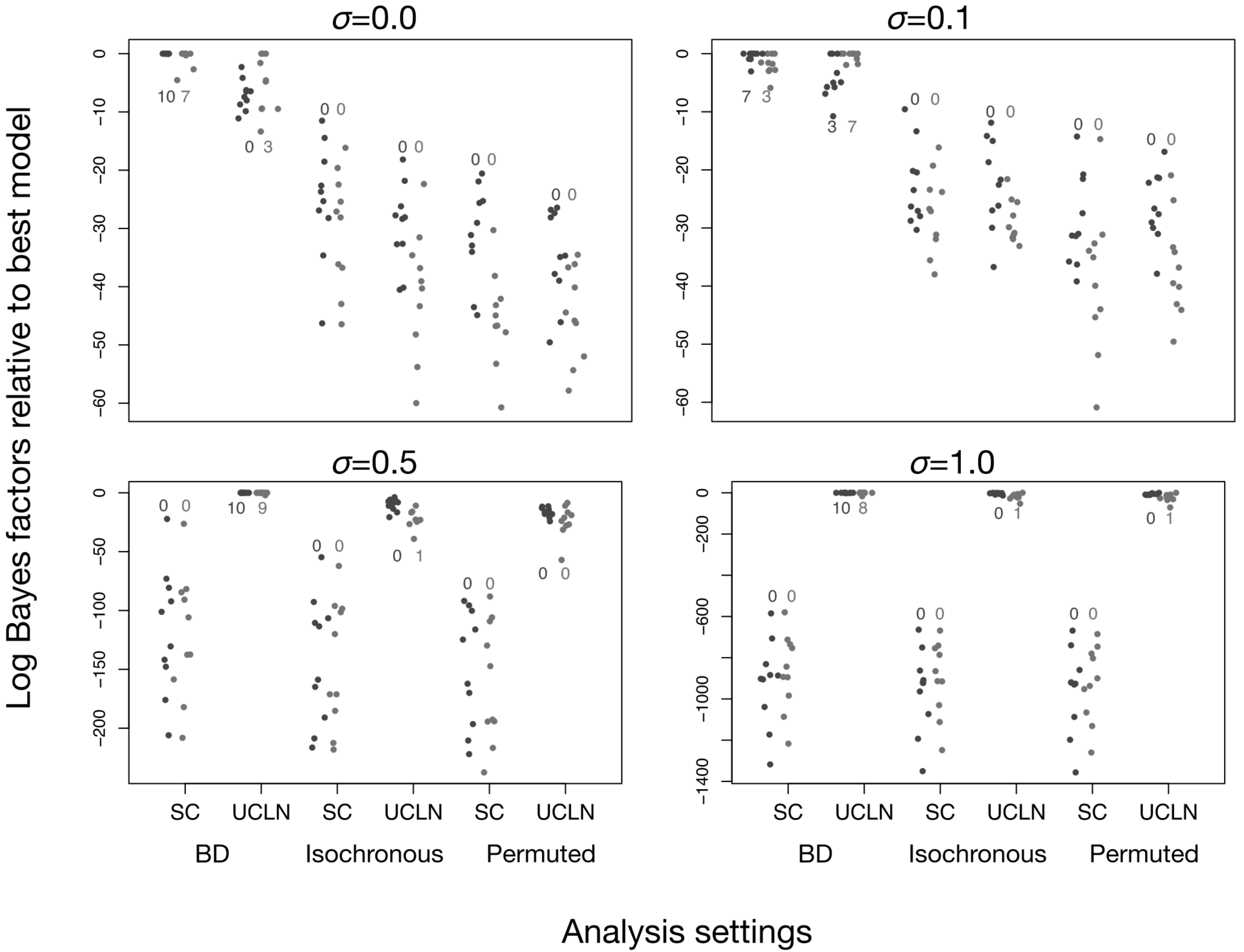

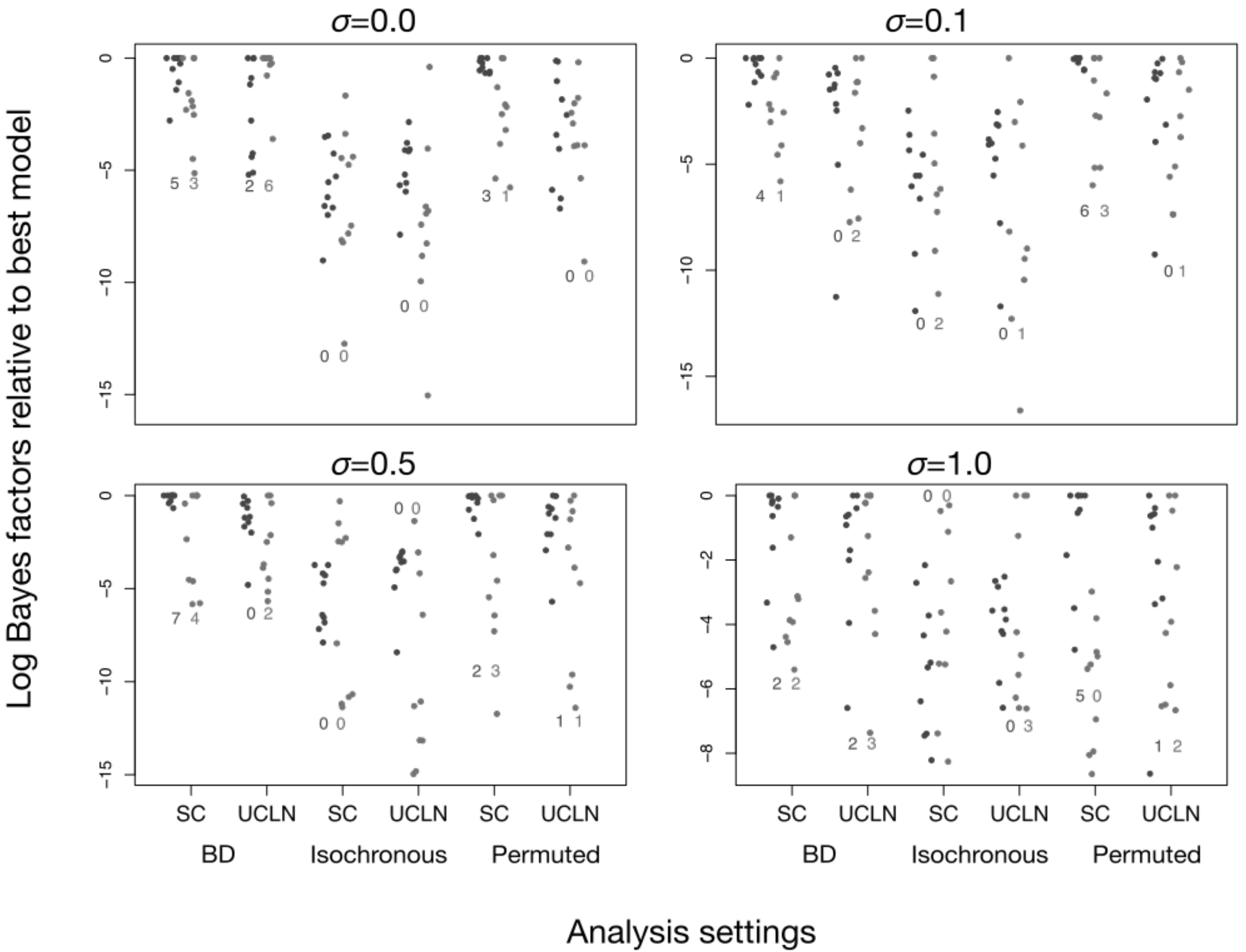

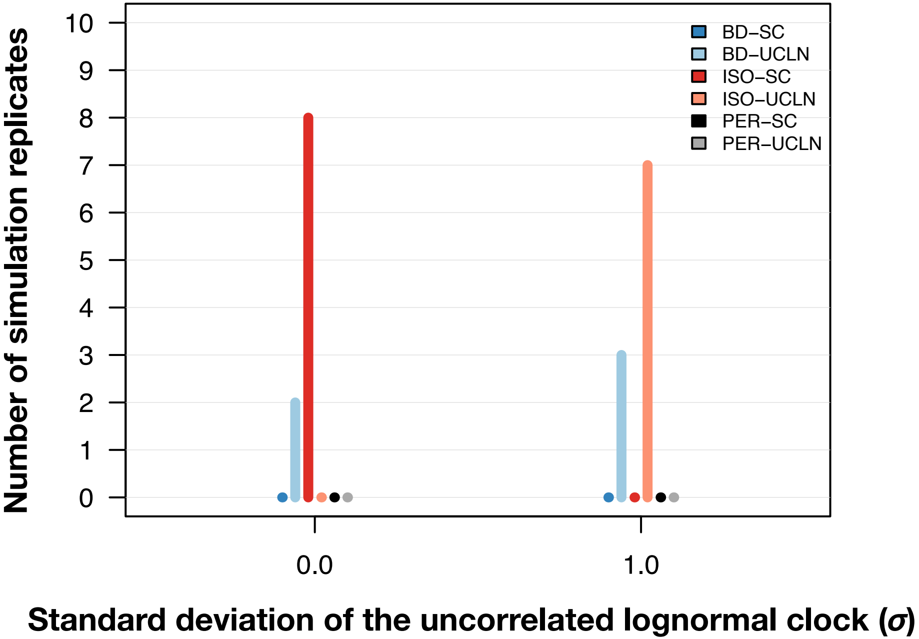
